# Limited metacognitive access to one’s own facial expressions

**DOI:** 10.1101/2021.03.08.434069

**Authors:** Anthony B Ciston, Carina Forster, Timothy R Brick, Simone Kühn, Julius Verrel, Elisa Filevich

**Author notes:** These authors contributed equally to the work (Anthony B Ciston, Carina Forster). **Corresponding author:** Elisa Filevich.

## Abstract

As humans we communicate important information through fine nuances in our facial expressions, but because conscious motor representations are noisy, we might not be able to report these fine but meaningful movements. Here we measured how much explicit metacognitive information young adults have about their own facial expressions. Participants imitated pictures of themselves making facial expressions and triggered a camera to take a picture of them while doing so. They then rated confidence (how well they thought they imitated each expression). We defined metacognitive access to facial expressions as the relationship between objective performance (how well the two pictures matched) and subjective confidence ratings. Metacognitive access to facial expressions was very poor when we considered all face features indiscriminately. Instead, machine learning analyses revealed that participants rated confidence based on idiosyncratic subsets of features. We conclude that metacognitive access to own facial expressions is partial, and surprisingly limited.

## Introduction

Precise motor planning and execution can occur without the brain having explicit, conscious access to the exact position of our limbs, or the exact degree of contraction of our muscles^1–3^. For instance, we can simultaneously walk, speak, and gesticulate successfully while concentrating on an argument and not on the movements that enable it, and we are furthermore unable to accurately report the state of each of our muscles. Although explicit access to proprioceptive signals in highly routinary tasks like walking or talking may be unnecessary, it might be beneficial in some other cases. For example, it has been suggested^4^ that metacognitive reasoning plays a central role in developing and improving motor expertise: if an experienced actor has a detailed and sophisticated representation of an ideal facial expression to communicate emotion, they are better able to detect and correct deviations from the ideal, leading in turn to more accurate and consistent performance.

Proprioceptive information about our limbs and their movements is thought to originate primarily from muscle spindles, together with skin receptors, Golgi tendon organs, and joint receptors^5–7^. Artificial vibration of the muscles can lead to activation of the muscle spindles, showing that their activation is sufficient to alter the representation of the body and its position^8, 9^. In addition, position estimates have been found to be more precise following active vs. passive movements, suggesting that efferent motor commands may either affect or inform proprioceptive representations^10–12^. Finally, proprioceptive information is combined with visual information, when available, to form a multisensory and integrated representation^13–17^.

Facial expressions present a particularly important yet poorly studied instance of motor control. On the one hand, we communicate a great deal of information with small, nuanced facial movements (on the order of 10 mm or less^18, 19^). On the other hand, we hardly ever see ourselves while making them. Perhaps with the exception of actors or public speakers who practice in front of a mirror (or the increased number of video-conferences during the 2020 SARS-CoV-2 pandemic), we do not usually have online visual feedback about our facial muscles. If visual feedback information is indeed critical to give rise to precise motor representations, facial movements might be very poorly represented. Together, the combination of the high social relevance of small movements in our facial muscles and the general lack of visual information about them raise the interesting question: How much do we know about how we look when we communicate with others?

Previous studies have focused on related questions. One line of research has quantified metacognitive access to *others’* facial expressions^20–22^ and operationalized metacognitive performance as the precision of participants’ representations of uncertainty. While our ability to accurately represent both the facial expressions of others and our certainty about them is clearly critical for social interactions, it is equally important to correctly represent and adequately control *one’s own* expressions^23^. In line with this notion, another line of research has aimed at measuring how accurate the representation of one’s own face is (under a neutral facial expression). One study^24^ found that participants showed a systematic bias to underestimate the length of their faces and slightly overestimate their width, mimicking what has been described for whole bodies^25^ and hands^26^. More recently, large inter-individual differences have been described in how accurately healthy young adults can represent their own faces^27^. These previous studies investigated relaxed faces with neutral expressions and captured, in essence, individuals’ ability to accurately describe their face, or to discriminate it from the face of another. Importantly, static features of one’s face are irrelevant to social interactions, which instead are based on dynamic information. Here, we focussed instead on metacognitive knowledge about how one’s face varies when making different expressions. In a pre-registered experiment, we asked participants to imitate expressions shown in pictures of themselves and to rate how well they thought they had imitated the expression. We then measured participants’ metacognitive access to their own facial expressions as the correspondence between subjective ratings and an objective measure of performance.

First, participants completed a task to measure their metacognitive access to facial expressions (Figure 1), consisting of three parts. Briefly, in the first part of the task, participants took pictures of themselves imitating different cue images done by actors^28^ to generate 32 participant-specific target images. In the second part, participants saw each of the target images on the screen and, while still looking directly into the digital camera, imitated themselves (Figure 1.B). In both the first and second parts of the task, participants pressed a keyboard key to trigger the digital camera. In the second part only, they additionally rated how confident they were in their own performance on a continuous confidence scale ranging from “Very unsure” to “Very sure”. Finally, in the third part of the task, participants saw the target and response pictures side-by-side and rated them for similarity on a continuous scale with the same labels as for the confidence rating. We quantified the distance between each image pair based on landmarks placed automatically on the pictures.

**Figure 1:**
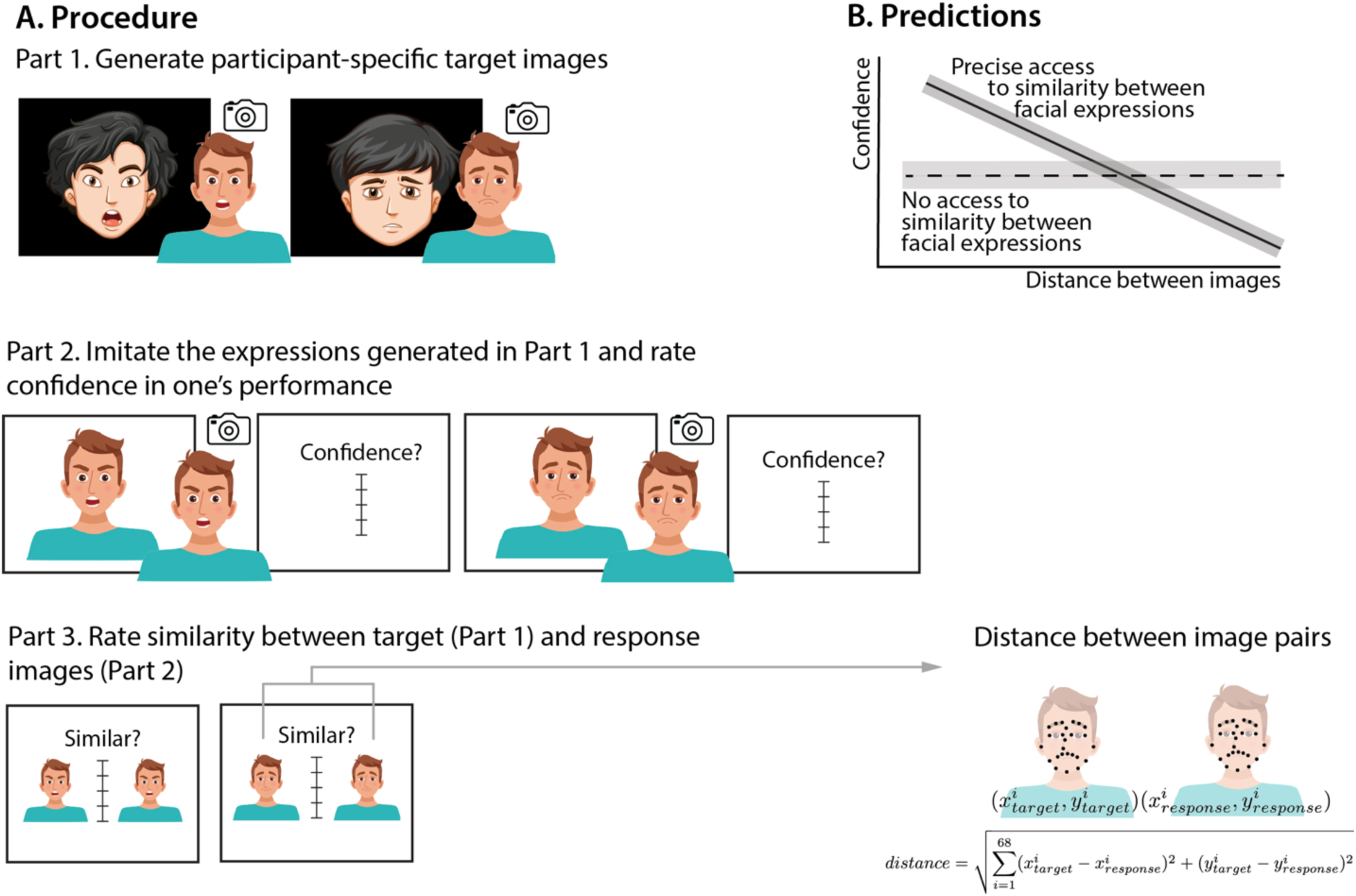
Experimental Design. **(A.) Procedure.** Cue stimuli were pictures of facial expressions taken from the MPI Small Facial Expression Database (Cunningham et al., 2005), but the images were replaced here with illustrations, to comply with the journal’s data privacy regulations. They were performed by actors and represented non-stereotypical expressions (e.g., “You lose the way in a foreign city”, see Methods for further details). Participants used these images as cues to produce 32 participant-specific target images. In part 2, each of the 32 target images (of the participants’ faces displaying the expression generated in part 1) was shown eight times (256 trials total). Participants reproduced their own expressions shown in the target pictures, pressed a key while holding their expression, and subsequently rated confidence in their own performance. The experiment was self-paced. Squares around the pictures indicate that they were displayed to participants, whereas pictures without a square frame around them represent pictures collected but not shown back to participants. (Expression drawing: Freepik.com) **(B.) Predictions.** The correlation between the two variables indicates the precision of the metacognitive representation. Confidence ratings were expected to be negatively correlated with the distance between two images if participants have metacognitive access to the low-level aspects of their facial expressions (solid line). Confidence ratings were not expected to vary with distance if participants had no metacognitive access to their own facial expressions (dashed line.

## Results

### Confirmatory Analyses

The distance between any pair of images is an inverse measure of performance in the task, as greater distance corresponds to a poorer match between target and response expressions. Thus, we reasoned that participants with precise metacognitive access to their facial expressions would have a sharp relationship between the distance between two images and the confidence ratings. The estimated regression coefficients from a multilevel model of these data should be negative and clearly different from 0. On the other hand, if a participant had no access to their own performance, their judgments would bear no relationship to the distance between two images, and the regression coefficients would be indistinguishable from 0 (Figure 1B, Predictions).

To arbitrate between these two possibilities, we first quantified our participants’ metacognitive access to their own facial expressions using a Bayesian linear mixed-effects regression model of participants’ confidence ratings. The model included the log-transformed distances as a fixed effect (for all 68 landmarks combined), as well as random intercepts for participant and facial expression. We found that participants’ confidence ratings had a small negative relationship to the distance measured (Figure 2.A, M = −0.03 ± 0.01, CI = [−0.05, −0.01], R^2^ = 0.21, see also Appendix 1-Figure 1 for the participant-wise data). However, when compared to the null model without the effect of distance, we found only anecdotal evidence^29^ for the relationship between the two (BF_10_= 2.20). Further, a robustness check revealed that, as expected given the proximity of the posterior samples to the region of practical equivalence (ROPE, defined following the default criterion of the region corresponding to a Cohen’s d of 0.1, Figure 2.B), the choice of the SD of the prior distribution had a strong effect on the BF_10_: Widening the prior distribution from 0.4 to 0.7 led to a BF_10_ = 1.02, and greater SDs also strongly reduced the value of the BF_10_. Together, these results point to no evidence for a relationship between confidence and distance. For illustration purposes, we plot the participant-wise posterior draws, in relationship to the ROPE (Figure 2.C).

**Figure 2.**
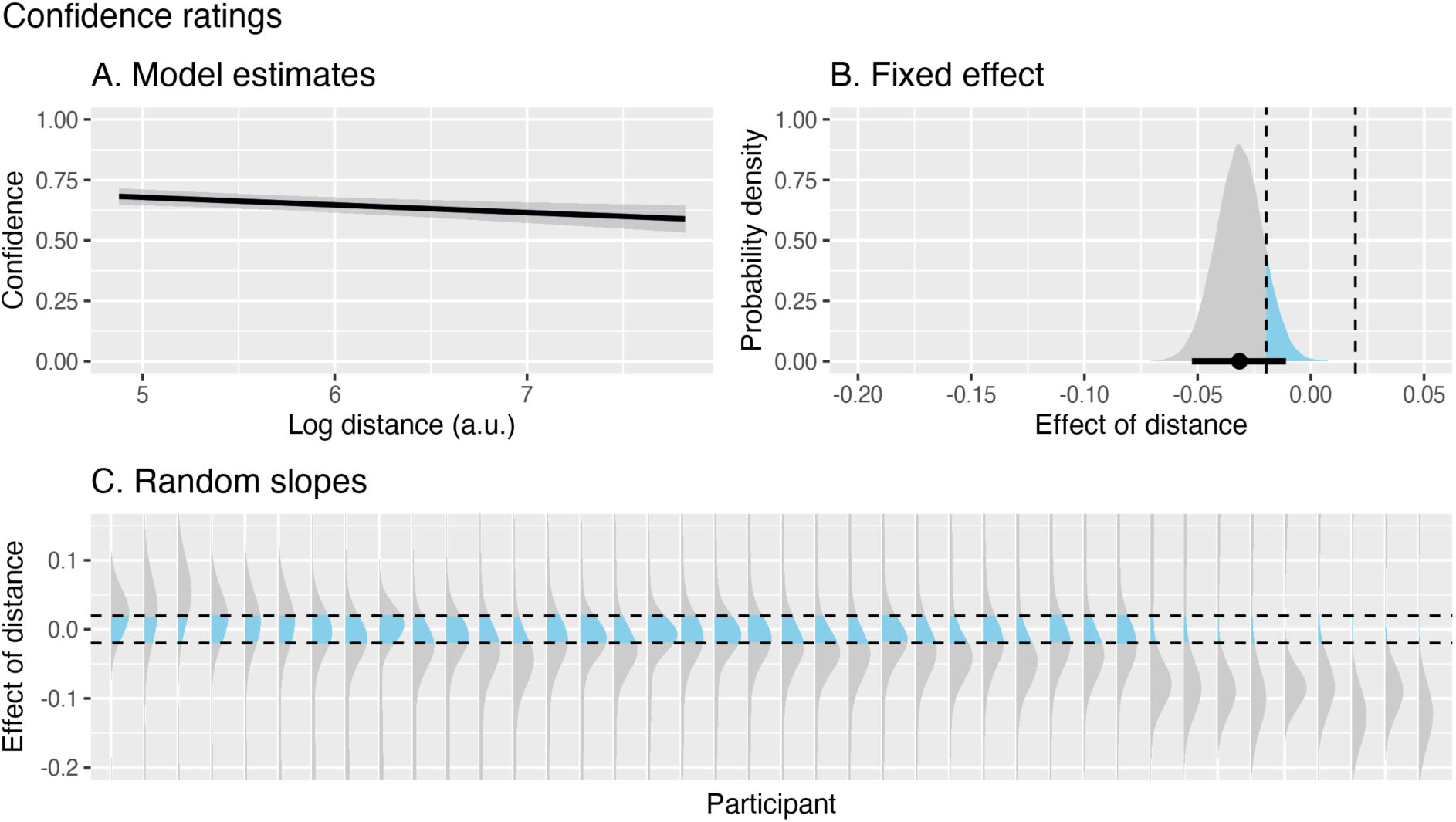
Poor metacognitive access to facial expressions. **(A.)** Group effects reflecting mean metacognitive access, namely the relationship between confidence ratings and distance between two images (inverse of performance). A small but consistently negative slope suggests that participants had minimal metacognitive access to their own expressions. The solid line represents the mean of the posterior draws, the shaded region represents the 95% credibility interval **(B.)** Posterior draws for the group-level fixed effect of distance, shown in relation to the ROPE, marked with dashed lines. The black horizontal line indicates the mean and 95% HDI. **(C.)** Posterior draws for each participant, shown in relationship to the ROPE. Note that the y-axis is clipped to better display the distributions around the ROPE and therefore excludes the long tails of some of the distributions. Participants are ordered following the mean slope estimate and might not be aligned across figures.

Then, to quantify the relationship between distance and similarity, we built a regression model of participants’ similarity ratings including, as before, the log-transformed landmark distances as a fixed effect (for all 68 landmarks combined), as well as random intercepts for participant and facial expression. Here, similarity ratings did track the distance (Figure 3 and Appendix 1-Figure 2). We found a clear and, as expected, negative relationship between the two (M = −0.12 ± 0.01, CI = [−0.14, −0.09], BF_10_ = 8.01×10^8^, R^2^ = 0.26). This shows that the distance we measured carried information relevant for similarity ratings and thus the null effect above cannot be simply due to a poor measure of distance. Additionally, because the same participants rated both confidence and similarity, the differences between the two ratings cannot be attributed to trivial effects such as a poor understanding of the confidence scale or task instructions, or simple lack of motivation.

**Figure 3.**
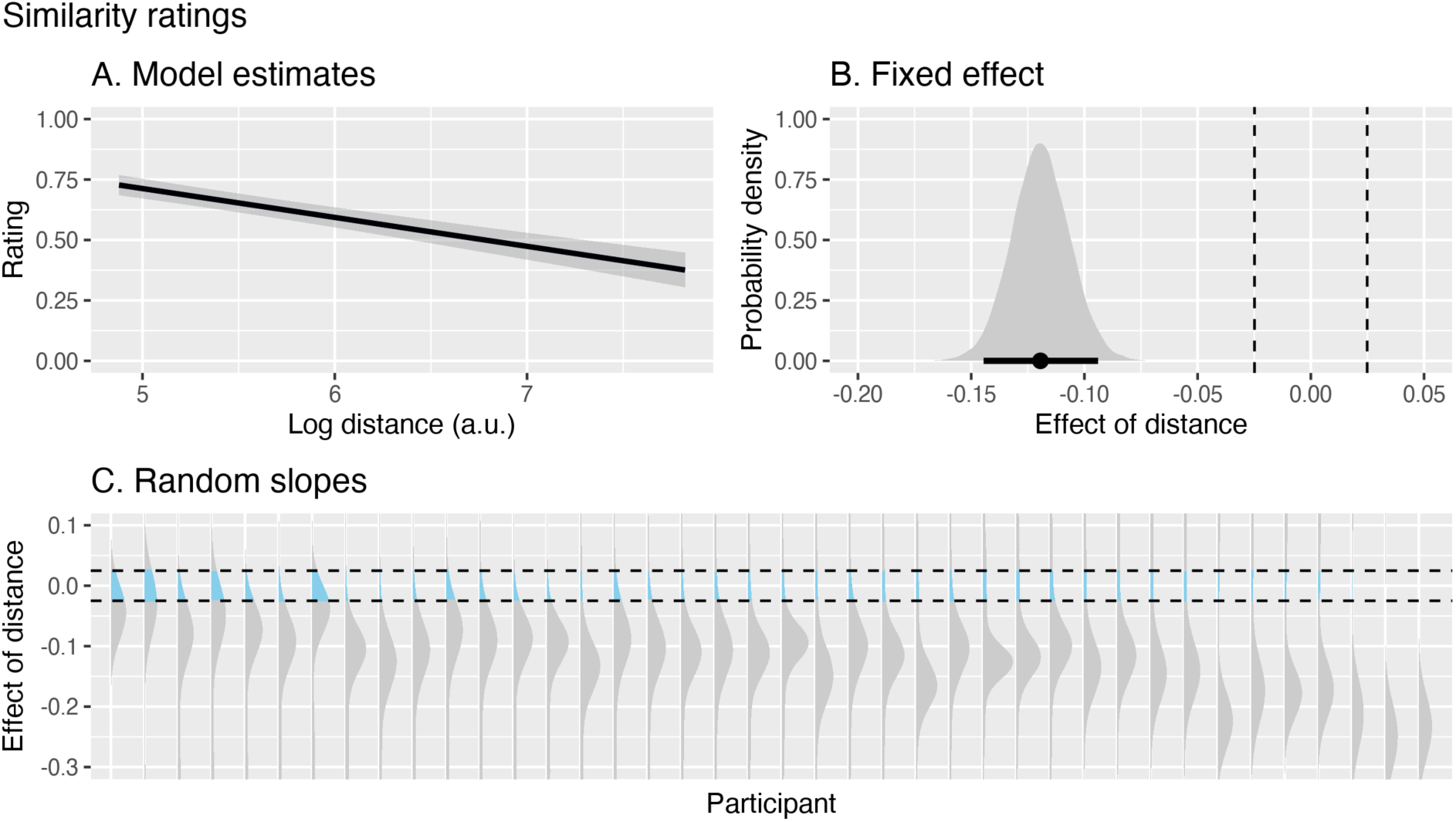
The distance between two images captures relevant information. **(A.)** Group effects reflecting the information contained in the distance between two images, namely the relationship between the similarity ratings provided by participants (when viewing each image pair side-by-side) and distance between two images. The solid line represents the mean of the posterior draws, and the shaded region represents the 95% credibility interval. **(B.)** Posterior draws for the group-level fixed effect of distance, shown in relation to the ROPE, marked with dashed lines. The black horizontal line indicates the mean and 95% HDI. **(C.)** Posterior draws for each participant, shown in relation to the ROPE. Note that the y-axis is clipped to better display the distributions around the ROPE and therefore excludes the long tails of some of the distributions. Participants are ordered following the mean slope estimate and might not be aligned across figures.

We emphasize that an advantage of similarity as compared to confidence ratings is almost trivial, as participants could see the picture pairs side-by-side to rate similarity, but not confidence. Hence, we simply take this result as a positive control to ensure that the landmark distances were at all related to similarity, but make no formal comparisons between the two kinds of ratings.

Finally, following our pre-registered plan, we explored relationships between the participant-wise random slopes with Mratio, a measure of visual metacognitive efficiency^30^ in a visual task. We found that visual Mratio was consistently above the chance level of 0 (M= 0.75, SD = 0.57, t(38) = 8.15, p < 0.001, BF_10_ = 1.54×10^7^, estimated with a default Cauchy prior) but that it did not correlate with participant-wise effects of distance on confidence (Figure 4.A, r = −0.19, p = 0.25, BF_10_ = 0.64, with a default shifted beta prior distribution). While the two measures of metacognitive access are not strictly comparable (the visual Mratio is controlled for first-order performance but the individual effects of distance on confidence are not), this analysis shows that poor metacognitive access to facial expressions cannot be attributed to generally poor domain-general metacognitive insight^31^.

**Figure 4:**
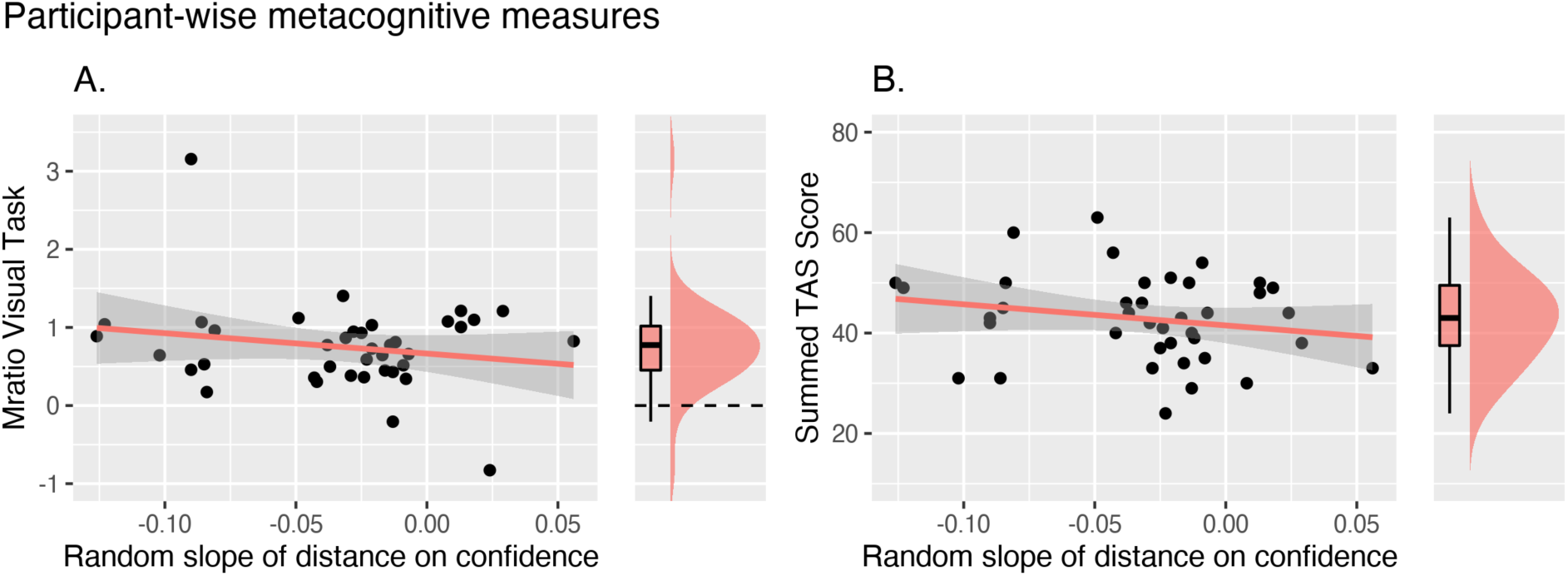
Correlations between participant-wise estimates of metacognitive access to facial expressions and other measures of insight. Each dot corresponds to one participant’s performance estimate, and the box- and density plots on the right represent the marginal distribution of the corresponding variable on the y axis. **A. Metacognitive efficiency (Mratio) in a visual task.** Participants’ metacognitive efficiency was significantly better than chance performance (marked with the horizontal dashed line). **B. Alexithymia score (TAS).** We found no evidence for a correlation between metacognitive estimates and these measures of insight.

Using Pearson correlations, we also measured potential associations between the inter-individual differences in metacognitive access to facial expressions and Alexithymia scores, as an indication of each participant’s ability to identify and describe their own feelings. We found no conclusive evidence for or against any relationships between alexithymia score and the participant-wise effect of distance on confidence (BF_10_ = 0.70, Figure 4.B) or on similarity ratings (BF_10_ = 0.43).

### Exploratory Analyses

For completeness, we studied the relationship between similarity and confidence ratings. We built a Bayesian linear regression model of participants’ confidence ratings, this time including the similarity ratings as a fixed effect and random intercepts for participant and facial expression. We found a clear positive relationship between the two ratings (M = 0.10 ± 0.01, CI = [0.09, 0.12], BF_10_ = 6.36 x 10^31^, R^2^ = 0.21, Figure 5 and Appendix 1-Figure 6). This suggests that participants’ confidence ratings were not random or noisy but rather that they simply did not reflect the low-level features captured by the distance.

**Figure 5:**
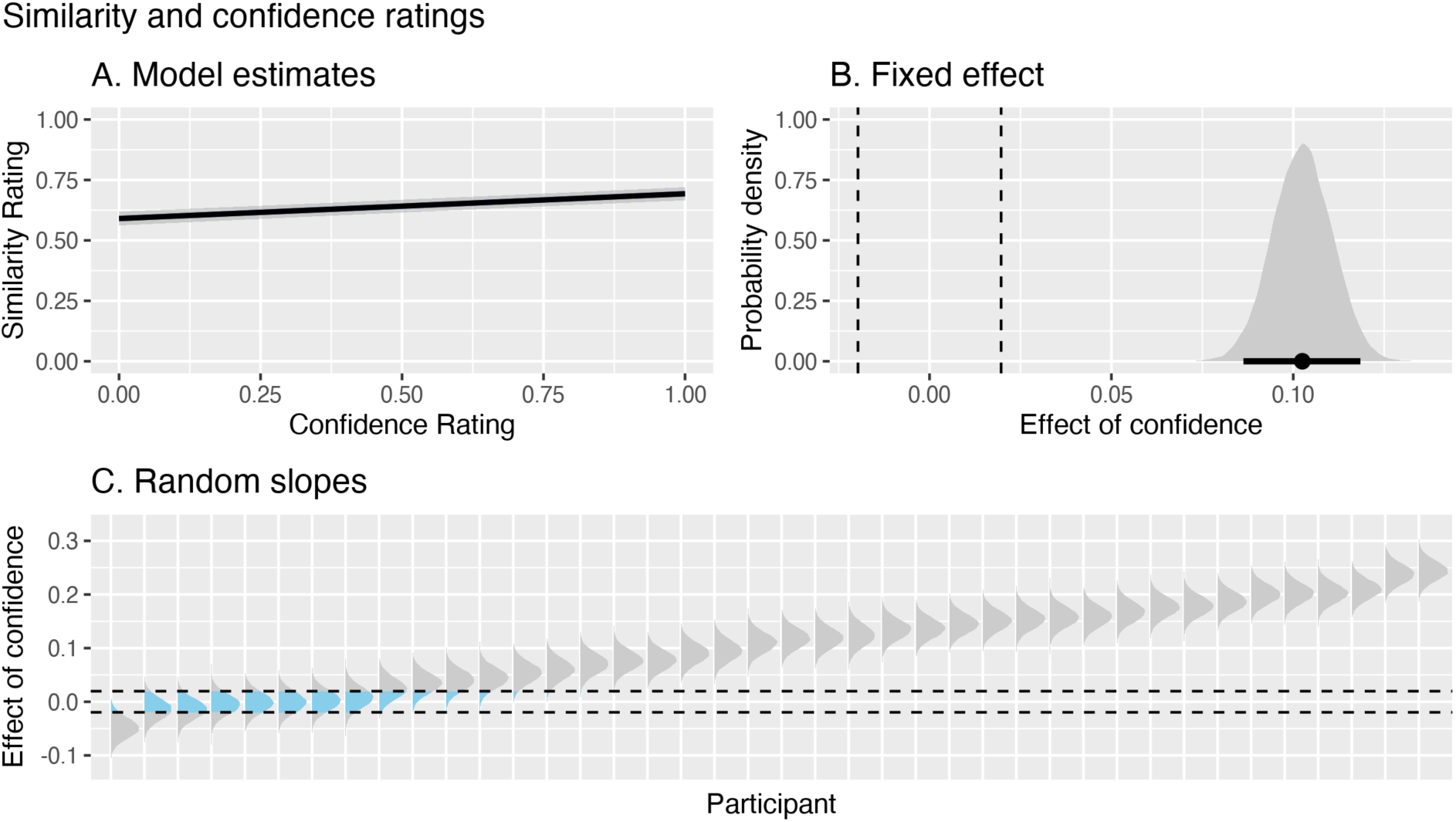
Similarity ratings vary with confidence ratings. **(A.)** Group effects showing the relationship between the two ratings on image pairs provided by participants (similarity vs. confidence). The solid line represents the mean of the posterior draws, and the shaded region represents the 95% credibility interval. **(B.)** Posterior draws for the group-level fixed effect of confidence on similarity, shown in relation to the ROPE, marked with dashed lines. The black horizontal line indicates the mean and 95% HDI. **(C.)** Posterior draws for each participant, shown in relation to the ROPE. Participants are ordered following the mean slope estimate and might not be aligned across figures.

**Figure 6:**
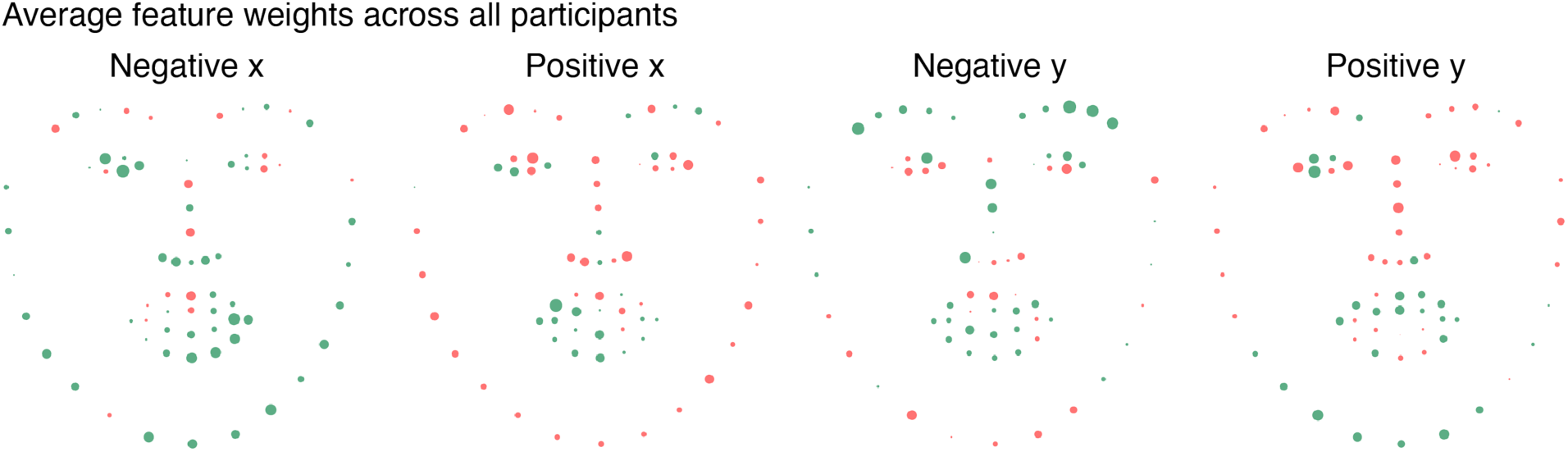
Machine Learning analyses. Average feature weights for participant-wise models of confidence ratings. Each dot represents the median feature weight for each landmark in models excluding RTs. Green and red correspond to positive and negative weights, respectively. The size of the dot corresponds to the relative magnitude of the landmark’s approximated weight within the model, and their positions correspond to a normalized face. Each landmark is split into the four cardinal directions, to yield four independent features (see Methods for details). We found no consistent pattern over participants where some features are weighted more strongly than others, see https://gitlab.com/elisa.filevich/cistonetal_metacognitionoffacialexpressions for an interactive table with participant-wise weights.

Our results so far suggest that participants’ confidence ratings did not reflect performance, calculated as the Euclidean distance over all landmarks. In a final set of exploratory analyses, we therefore aimed at identifying which pieces of information participants may have taken into account when rating confidence.

The Euclidean distance between image pairs assigns equal weights to the distances of all facial landmarks and is therefore a relatively naive measure of the difference between expressions, in that it does not allow for potential differences between landmarks in their contribution to different individuals’ confidence. However, it is in principle possible that participants attended to different parts of their faces to different degrees and, further, that this differential attention was not consistent across participants. For example, one participant may have focused almost exclusively on how well their mouth matched the target image to rate their confidence, and another participant may have focused exclusively on the eyes and ignored the mouth. While this was against the task instructions, it remains a possibility that would undermine the strong claim that most participants did not base their confidence ratings on the landmark distances. To obtain a more fine-grained and flexible measure of performance we used a simple linear regression machine learning (ML) model to predict each participant’s confidence ratings using a principal component (PC) decomposition of the distances between corresponding landmarks as features. Building participant-wise models provided the maximum flexibility in feature weight assignment and was therefore the harshest test to the conclusion that metacognitive access to facial expressions is poor. We found that these models could in fact predict confidence ratings (median *r* = 0.26 ± 0.15), suggesting that participants did indeed base their confidence ratings on (specific subsets of) landmark distances. Further, because confidence is known to correlate negatively with response times^32, 33^, we also asked whether RTs could have served as a proxy for distance. We found that the landmark distances could be used to build ML models that predicted confidence ratings above and beyond RT information alone, confirming that participants did use some of the landmark distance information to rate confidence (see Appendix 1-Figure 4).

To better understand which information participants used to rate their own performance, we reconstructed the weights of each feature in landmark space (based on the model’s weighting of each principal component and each feature’s loading on that component, see Methods). We first plotted the resulting landmark weights on their corresponding mean locations to explore potential patterns among participants based on the set of landmarks with the highest weights (both visually and by considering the median weight over all landmarks); however, we could not identify any landmarks or features that were consistently prioritized across participants (Figure 6). Individual participants’ ML feature weights can be seen at https://gitlab.com/elisa.filevich/cistonetal_metacognitionoffacialexpressions/). Finally, we estimated the relationship between the new landmark distance (this time considering the participant-specific weights) and confidence ratings using, as before, a linear mixed-effects regression model. In line with the non-zero r values from the ML models, the reconstructed distances did in fact show a significant relationship with confidence ratings (M = 0.04 ± 0.004, CI = [0.03, 0.04], BF_10_ = 1.34 x 10^7^, R^2^ = 0.24). Note that the slope estimate is now positive, because the feature weights must incorporate the negative relationship between landmarks and confidence, in order to predict confidence ratings. Taken together, the results suggest that participants were indeed able to base their confidence ratings on the distances between facial landmarks, but only on a subset of them; and that each participant had access to, or focused on, different aspects of their facial expressions.

## Discussion

We asked how much we know about how our faces look when we make expressions. We quantified young, healthy adults’ metacognitive access to the low-level details of their own facial expressions. We emphasized to participants that we were focused on the specific shape of the face and activation of the muscles, not on the emotion that the expression conveyed. Surprisingly, our results suggest that participants were only very poorly able to consistently base their confidence ratings on the complete set of facial features. A priori, this can be interpreted in two (non-exclusive) ways: Participants’ confidence ratings may not have strongly relied on the distance between a pair of images because they truly had little or no metacognitive access to their own facial expressions. Alternatively, our measured distance based on the whole set of landmarks may have been a very noisy or even invalid measure of performance. In turn, this alternative explanation would mean that it would be invalid to quantify metacognitive access as we did. To ensure that the second alternative could not fully explain our results, we quantified the relationship between ratings of similarity (provided by the participants themselves while viewing image pairs side-by-side) and distance (based on the whole set of landmarks, combined with equal weights). Here, we did find a clear relationship between the two, suggesting that the distance between image pairs does carry information that is — to some extent — relevant for similarity. This result also shows that a poor relationship between confidence and distance cannot be attributed simply to poor use or understanding of the confidence scale. It is important to emphasize that we draw no conclusions from the direct comparison of the strengths of the association between distance and the two kinds of ratings (namely confidence and similarity), as it would not be a valid comparison. Participants had no visual information about the expression they were making when rating confidence, whereas they could do careful comparisons of image pairs using all available visual information to rate similarity. Instead, we make separate inferences based solely on the estimation of the effect size and reliability for each of the associations, and the comparison between each full model including the effect of interest and its null counterpart. Simply put, the analysis of the relationships between confidence and distance suggests that participants could access their performance only poorly. On the other hand, the analysis of the relationships between similarity and distance suggests that we measured performance adequately.

Beyond the group-level effects, we found variation between individuals. We aimed at explaining this variation by exploring correlations between these individual estimates of the relationship between distance and confidence and other measures of insight, namely visual metacognitive efficiency and alexithymia score. No conclusive relationships emerged that could explain the variations between individuals.

Further, in another exploratory analysis, we considered that the summary distance measure could not discriminate between landmarks that heavily informed participants’ confidence ratings and those that were ignored. In other words, confidence ratings may have depended on performance defined by a subset of landmarks, which may not have been the same for all participants. To examine this possibility, we built linear regression ML models on confidence ratings that included the differences for each landmark as individual features (each of them separated into the four cardinal directions). This analysis revealed that the models built for all participants could predict confidence from the combined features (and could do so with better accuracy than the models relying solely on reaction times, which we expected to be predictive of confidence based on previous literature^32, 33^). This result suggests that participants’ confidence ratings do indeed carry information about the landmark distance between target and response expressions. But, unlike what the linear regression analyses assumed, not all landmarks contribute equally. In fact, some landmarks contributed in a way that was contrary to what was expected (i.e. larger distances were associated with higher confidence). Further, the contributions from each landmark were not consistent between participants. In sum, because some variability in facial expressions did not appear to inform confidence ratings, we argue that these findings show that there is a disconnect between participants’ ability to control their faces (through their low level features) and their assessment of performance. While some aspects of participants’ facial expressions led (idiosyncratically) to higher confidence ratings, these ratings were not indicative of performance.

If it is indeed the case that young, healthy volunteers have only partial access to their own facial expressions, the obvious question arises: How do we communicate effectively in society? Drawing from previous literature, we assume that each facial expression carries both low-level information (the specific degree of contraction of each muscle and consequent location of the landmarks) and high-level information (the emotion conveyed) and that these two bits of information are not necessarily correlated. We note that the effects we observed here are valid for the low-level features which we asked participants to concentrate on, but they may not extrapolate to the high-level features of facial expressions.

In fact, we suggest a simple model (Figure 7) consistent with our results where these two aspects are dissociated. We obtained the distance using an algorithm that, we assume, has no access to high-level information. Similarity ratings, on the other hand, were made by human observers (the study participants) and therefore were based on both the low-level features (by design, in line with our instructions) and high-level emotional information that is automatically processed^34^, as we discussed above. On the basis of our results, we contend that confidence ratings may be based chiefly on high-level information, as they can only poorly incorporate low-level information. Then, the shared (high-level) information between similarity and confidence ratings explains why they correlate and the dissociation between low- and high-level information, together with their unequal contribution to different ratings, explains why confidence and distance are in turn dissociated.

**Figure 7:**
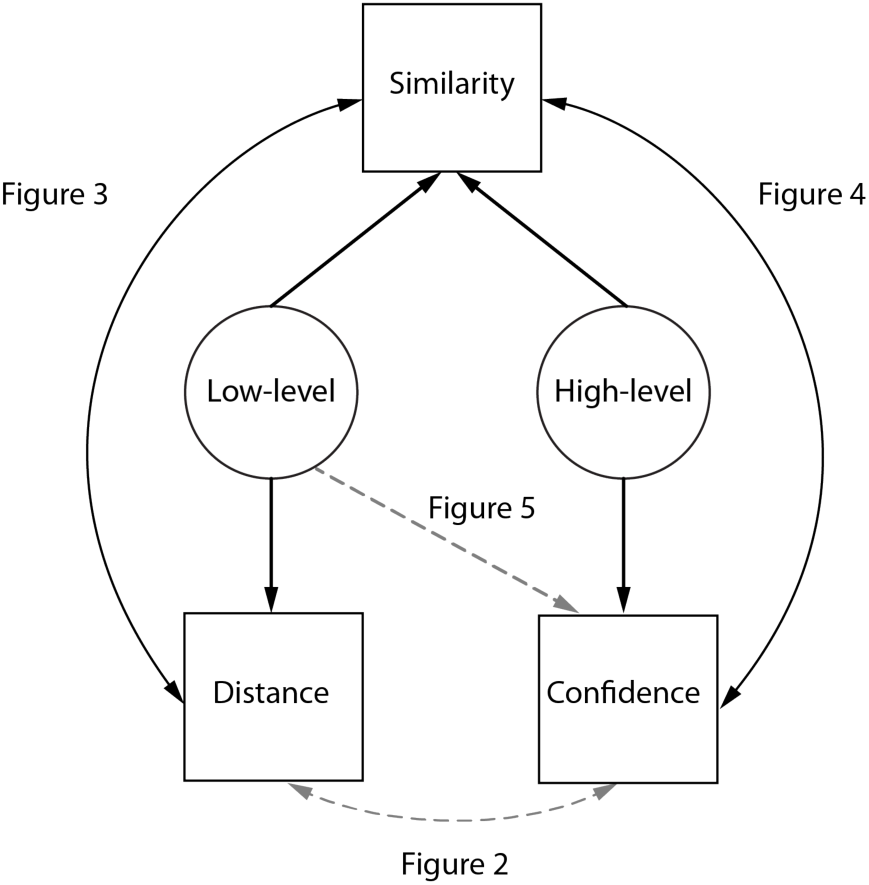
Suggested model for metacognitive access to facial expressions. We consider that each facial expression carries both low-level and high-level information (here depicted as circles because they are akin to latent variables in a structural equation model, whereas the measured variables of Distance and Confidence are depicted as squares). We also consider that the distance we measured is solely based on low-level information that the algorithm has access to. Thus, this simple suggested model (where confidence has accurate access to high-level but poor or partial access to low-level information, and where similarity ratings by human judges are informed by both low- and high-level aspects of each image) is sufficient to explain both, on the one hand, the relationships that we observed between distance and similarity and between similarity and confidence, and on the other hand, the dissociations we found between confidence and distance.

The distinction between metacognitive access to high- and low-level features of facial expressions is compatible with previous literature. It has been shown that the brain regions involved in assigning confidence to the accuracy of purely perceptual decisions (the thickness of a horizontal bar presented above-fixation) were different from those assigning confidence to decisions about emotional faces^20^. Two recent studies presented participants with two conditions with more closely matched stimuli. In the first one, two groups of participants underwent one of two kinds of perceptual learning^21^. One group trained to discriminate between two faces based either on their identity (high-level features) and the other group trained to discriminate the contrast between two faces (low-level features). The results showed that, while there was perceptual learning (first-order performance remained stable despite increased task difficulty) in both groups, metacognitive accuracy improved for the low-, but not high-level features training group. The authors argued for a dissociation between metacognitive access to these two levels and for a dual-stage model of metacognition whereby perceptual learning reduces noise in the representations for low- (but not high-) level facial features. A second study used a causal intervention^22^ to show that continuous theta-burst suppression to the lateral prefrontal cortex led to a decrease in metacognitive performance in a task that relied on the low-level aspects of faces (discriminating between the orientation of two faces) but not one that relied on high-level aspects (discriminating the expression they communicated). Together, these results support a distinction between metacognitive access to high- and low-level features of *seen* faces (i.e., others’ faces). We extend these results and suggest that this distinction may also apply to the case of one’s own face, even when not seen.

Facial muscles appear to lack muscle spindles^35–38^, which are the main sensors for skeletal muscle stretching^5–7^. Instead, other mechanoreceptors have been suggested to replace muscle spindles in their transduction of electric signals elicited by facial muscles^39^. In contrast to what we described for facial muscles, young, healthy participants have above-chance and precise metacognitive access to movements that are controlled by skeletal muscles^40^. Moreover, unlike the case of metacognition of facial expressions, measures of metacognitive performance in motor control do partially correlate with those from a visual task^41^. Speculatively, at least two factors may explain these discrepancies. First, different stretch receptors may lead to different kinds of representations that may be differentially accessible to metacognitive monitoring. Second, visual feedback during development and motor learning might play an important role. Extensive motor learning and concomitant visual information for limbs that are in the field of view may shape and lead to sharper conscious representations in a way that is not possible for facial expressions.

### Relationship to other metacognitive tasks

Many of the recent studies measuring metacognitive performance have capitalized on a relatively rigid operationalization of metacognition that quantifies metacognitive performance as the relationship between subjective confidence ratings (the second-order task) and objective performance in a 2AFC (the first-order task), and especially in whether a participant is able to assign high confidence exclusively to correct trials^42^. Unlike most experiments on metacognition, where experimenters can very easily control the (often visual) stimuli that they present to participants, the study of motor metacognition requires participants to make a movement in the first place, thereby adding another task to the standard operationalization. Participants make a movement (zero-order), then make a (first-order) judgment about it, and finally provide a (second-order) subjective confidence rating. Examples of a zero-order task include moving a finger at a given pace^40^ or throwing a ball to hit a target^41^. A different approach, which we took here, consists in operationalizing the metacognitive judgment not as confidence in accuracy of a binary choice, but instead as a judgment of performance^43–45^. While both operationalizations may be valid, it is important to note the differences between them to prevent assuming unwarranted relationships: The first approach, borrowed from paradigms developed for perceptual tasks, makes a very clear distinction between three different tasks with, in principle, independent performance levels. In a ball-throwing task, a person could miss a target often (poor zero-order performance), be good at discriminating whether the movement they made would hit the target or not (high first-order performance), but assign high and low confidence equally often to correct and incorrect discrimination trials (low second-order performance). This sharp distinction between three cognitive levels is elegant and makes metacognitive motor tasks directly comparable to perceptual ones. On the other hand, the comparison may not be as straightforward as it appears to be^46^. It has been argued that this rigid operationalization ignores a distinctive feature of (sensori)motor performance monitoring: In making a movement, we must monitor our performance in relationship to the intended goal, which includes not only perceptual uncertainty but also motor noise and skill^43, 47^. Thus, the approach of asking participants to rate their own performance allowed us to measure metacognitive access as the relationship between true performance and the (arguably) ecologically relevant estimate of subjective performance.

### Introspective vs. extrospective access

These results contribute with an interesting case to the question of introspective privilege. A classic view has argued that introspection has privileged first-person access to — and is thus the ultimate authority on — mental and emotional states^48^. In the motor domain, this would mean that the agents always have the most precise representation of their movement. This makes intuitive sense, as a precise representation of an ongoing movement is presumably a prerequisite for fine and efficient motor control and execution, as well as for the emergence of a sense of agency^49, 50^. On the other hand, a reading of the empirical literature does not provide a clear answer, perhaps due to the diversity of motor paradigms examined. Some studies have shown that precise access to movements is not always available at an explicit representational level. Participants failed to report large corrections to their ongoing movements^51^, and explicit instructions about how to solve a visuomotor rotation task can in fact be detrimental for performance, because explicit control is not a substitute for implicit corrections, which occur without participants’ awareness^52^. Healthy participants also appear to have poor access to their own eye movements and a poor (i.e., noisy) representation of their own bodies that can be easily affected by visual cues^13, 14^. On the other hand, almost directly contradicting the results above, other studies have shown that metacognitive representations of movements are as precise as those of exteroceptive signals^40^ and that explicit instructions can sometimes be indeed beneficial for performance by leading to quicker adaptation times and shorter after-effects, as compared to no explicit instructions^53^. To understand these discrepancies, it may be helpful to measure metacognitive access systematically across different muscle effectors and motor and metacognitive tasks. By examining healthy participants’ explicit knowledge of their own facial expressions, then, we explored another — and in our view very important — instance of motor control. We suggest that, perhaps just like eye movements, some parts of motor control might be opaque to explicit introspective access. This contributes to the body of literature questioning the privileges that introspective access has been argued to have as a matter of principle and levels the balance of epistemic access towards the complementary notion of extrospection^48, 54^.

### Limitations

One important limitation in our analyses is related to one basic assumption of our approach. In our exploratory analyses, we found a clear relationship between confidence and similarity ratings at the single-participant level. We explicitly relied on the distance estimated by the algorithms as the ‘true’ measure of performance. We argue that this assumption is valid for two main reasons. First, we specifically instructed participants to focus on these low-level aspects. Second, we found very similar results using two completely different algorithms to place facial landmarks (see SI), suggesting that this measure of distance captures true differences in facial features and does not depend heavily on the idiosyncrasies of the algorithm. However, it could be argued that similarity ratings are in fact a better, truer measure of performance because they reflect how similarly two faces are perceived by a person (either a judge or the very same participant) in an ecologically valid setting. Against this intuition, we argue that similarity ratings could have been subject to the same biases and heuristics that confidence may have relied on. As a very simplistic example, a given participant could have consistently rated positive expressions with higher confidence and similarity than negative expressions, leading to a relationship between the two kinds of ratings that needn’t be explained by metacognitive access. We note, however, that this alternative analysis of the data, based on different assumptions, would have led to the cardinally opposite conclusion that participants *do* have precise metacognitive access to their own expressions.

A second limitation has to do with the predictive power of our statistical models. Despite robust effects in the Bayesian mixed models, a significant amount of variability is left unexplained (see SI). Better measures of distance, more precise motion tracking technologies (like infrared reflectors placed on the face), or different analysis methods may have reduced this unexplained variance. Additionally, we note that our analyses are based on static images, namely the endpoints of otherwise dynamic expressions. But, important information is conveyed in the dynamic pattern of facial expressions^55–57^, and a future direction of this work might be to relate confidence to dynamic aspects of facial expressions instead.

Finally, while the exploratory machine learning analyses allowed us to identify potential aspects of the face that participants attended to while ignoring others, we might have failed to detect any true effects where the relationship between confidence and distance differed between expressions, or relationships that changed significantly over the course of the experimental session.

It could be argued that the use of non-canonical expressions limits the ecological validity of our paradigm. However, we note that in this study we were interested in studying a potential disconnect between (zero-order) motor control and (second-order) metacognitive access to it. Canonical expressions, where a highly trained and stereotypical set of movements correspond, one-to-one, to a specific expression, confound motor control with emotional content and would not have allowed us to make any inferences about which kind of information participants were accessing to make their judgments. For instance, had we asked participants to make a stereotypical “happy” expression and then rated confidence, we would not have been able to determine whether their confidence judgments were well calibrated with the emotional state they recreated, the highly-trained motor program, or the end state of the target expression. In short, canonical expressions would have carried with them a set of confounds that our paradigm avoided.

## Conclusion

Our analyses suggest that healthy young volunteers were only able to estimate their performance in producing non-stereotypical facial expressions based on partial information. This indicates that we not only do not have metacognitive access to the low-level details of our facial expressions, but also suggest that we cannot access them, even when explicitly asked to do so under experimental conditions. This is surprising, we argue, because it sets facial movements apart from other body movements (namely those of arms and fingers), for which, as previous studies have shown, we do have precise metacognitive access to lower-level motor information, even when this information is decoupled from the motor goal. We speculate that this distinction might be related to the lack of concurrent visual information during social interactions, but our speculation will need to be examined in future studies.

## Material and Methods

### Participants

Following our pre-registered plan (https://osf.io/pnyw3), 40 healthy participants took part in the study after giving informed consent (21 female, 19 male mean ± SD: 28.2 ± 4.6 years). We based the sample size on pilot data from 12 participants (see SI) and previous studies of motor metacognition from our group. Exclusion criteria were a recent history of psychiatric disease or having a heavy beard, as we reasoned that it would occlude the view of part of the face and placing of the landmarks. The local ethics committee approved all procedures (Nr. 2017-23-R), which conformed to the Declaration of Helsinki.

### Apparatus

The experimental setup consisted of a stimulus computer, a digital camera, a screen, and a half-silvered mirror tilted 45° from the vertical (Figure 8). Participants saw the image displayed on the screen by the stimulus computer indirectly through its reflection on the half-silvered mirror. Behind the mirror, a digital camera (Fire-i, UniBrain, Athens, Greece) connected to the computer took pictures of the participants’ facial expressions. This setup allowed participants to look at the pictures displayed while simultaneously looking directly into the camera. As a result, we obtained pictures of participants looking straight ahead and not downwards at the image, as would have been the case if we had used e.g. a simple laptop computer with a digital camera just above the screen.

**Figure 8.**
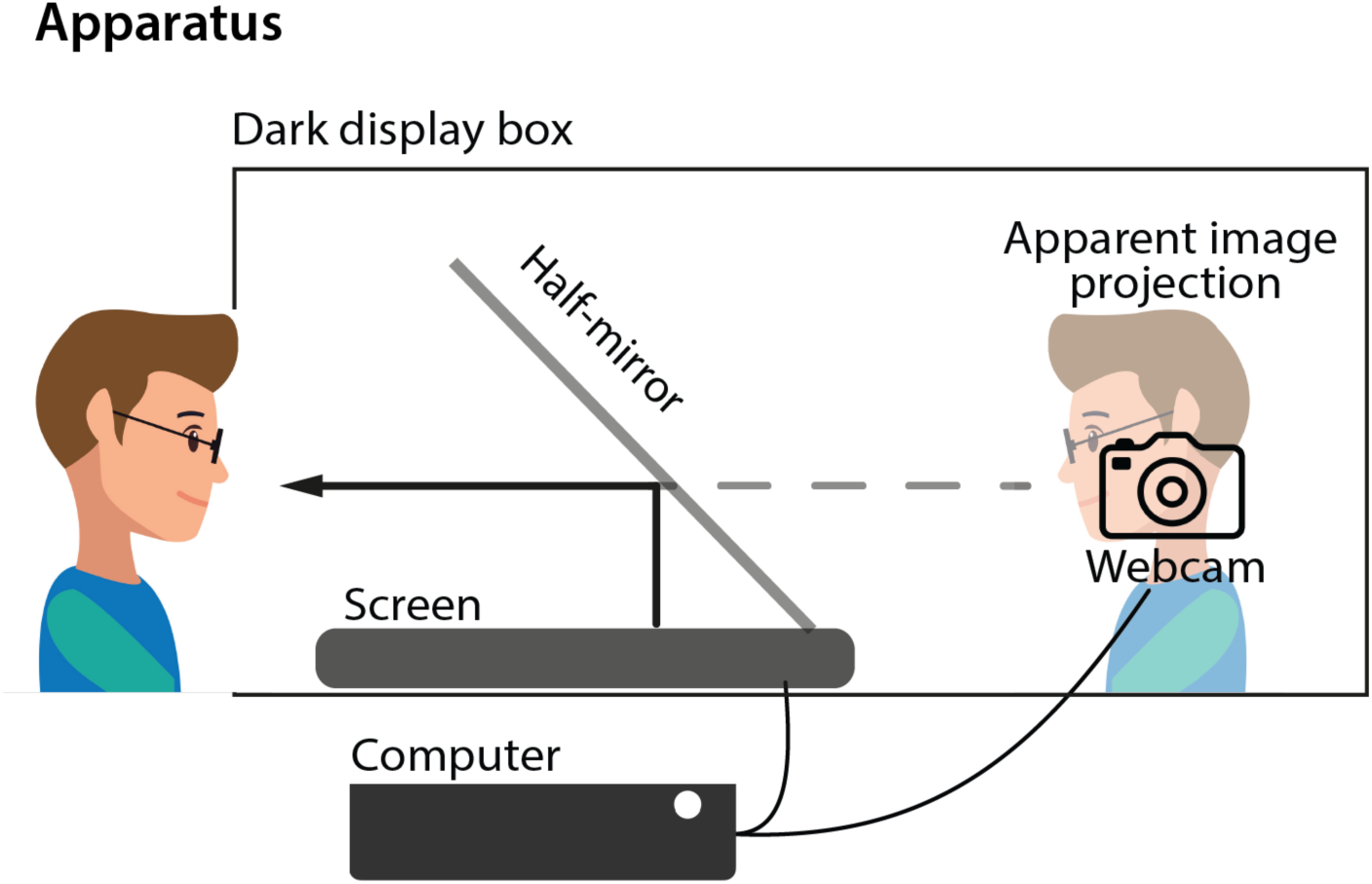
Experimental Apparatus. Participants sat in front of a dark display box and saw the pictures projected from a computer screen reflected on a half-plated mirror (tilted 45°). Behind the mirror, positioned directly in front of participants’ gaze, a digital camera took pictures of the participants when they pressed the corresponding key. This way, participants could look simultaneously directly at the to-be-imitated picture and into the camera.

Participants sat at approximately 60 cm from the middle-point of the half mirror, which was in turn 45 cm away from the display screen. In order to reduce head movements, we held participants’ torsos loosely in place with an elastic band tied to the chair. Additionally, at the beginning of the experiment, we showed participants the image collected by the camera in real time and asked them not to make large head movements or rotations. While it would have been desirable to further limit whole-head movements using, e.g., a chin rest, we opted against this as it would have made expressions unnatural and, more importantly, because it would have provided a form of sensory feedback, interfering with the experimental design. We ensured that participants’ faces were well-lit and took care that participants did not see any reflections of their own face on the mirror.

### Procedure

All experimental tasks were written on MATLAB (R2016b, The Mathworks, Natick, MA), using Psychtoolbox-3^58–60^ and ran on MacOS. All tasks were self-paced with no time deadlines. All participants (except for one, due to technical problems) completed all tasks in the same order.

### Facial Expressions Task

The facial expressions task consisted of three parts. In the first part (Figure 1.A), participants saw 32 different pictures of four different actors in pseudorandomized order (see the description of *Cue images*, below) and imitated each expression as best they could. Participants pressed a key (the space bar) once they considered that their expression was as close as possible to the actor’s expression. We asked participants to try to match the low-level physical features of the face — the curvature of the lips, the elevation of the eyebrows — rather than the emotion conveyed by the expression. Upon pressing the spacebar, the digital camera behind the half-plated mirror took a picture of the participant’s facial expression, and a new trial started. On a separate test, we had determined that there was a minimum delay of approximately 80 ms between the time of key press and the time stamp of the image. Accordingly, we included in our instructions to participants to hold the expression in place after they had pressed the key that would trigger the image acquisition.

The 32 pictures of participants generated in this way served as target images for the second part of the paradigm. Here, participants saw the target images and tried to reproduce their own expressions. Once again, we emphasized that the goal was to match the low-level physical features of the face rather than the emotion conveyed. After each trial, participants used a mouse to rate their confidence (on a visual analog scale) regarding how well they thought that they had imitated their own previous expression. Participants saw each of their 32 target expressions repeated 8 times in random order (256 trials in total). We only revealed that they would have to reproduce their own expressions after the first part of the experiment was complete. Parts 1 and 2 of the experiment took on average approximately 50 minutes. Before starting part 1, participants completed four practice trials where they simply imitated pictures of famous celebrities and took pictures. They did not see the resulting pictures of themselves.

In the third part of the task, participants saw each of the 256 pairs of pictures (target and response) and rated them for similarity on a scale exactly like the one they had used for confidence. This part of the experiment took on average 30 minutes.

### Cue images

We used 32 different facial expressions as cue pictures (14 from two different male actors, 18 from three different female actors) which would be used to generate participant-specific target expressions. To prevent participants from producing stereotypical target expressions, we sought pictures representing expressions that could not be unambiguously categorized as one of the basic emotions^61^. We selected pictures from the MPI Small Facial Expression Database^28^, which includes video sequences of expressions based on a method acting protocol in which actors produce non-standard expressions by imagining themselves in a situation described by a brief scenario and reacting accordingly. Example descriptions of expressions include: “Somebody suggests to try something. You hesitate at first, then you agree”, or “You have reached a goal and you are happy to have accomplished it”. Additionally, we selected still images from the video sequence that did not correspond to the peak expression, but instead to an intermediate step. As a result, the cue images could not easily be labeled as stereotypical expressions (e.g., “happy”, “sad”) for which participants might have a predefined motor program but could instead be assumed to be the result of an unusual and idiosyncratic combination of gestures. Note that, as the samples in Figure 1.C show, these cue images were not unnatural grimaces and so the paradigm remains ecologically valid. We reasoned that these non-canonical expressions would maximize motor variability, ensuring that confidence ratings could be based only on a true evaluation of trial-by-trial performance and not on a general knowledge of how reproducible a given expression was.

### Visual Task

Each participant completed 200 trials of a visual metacognition task (https://github.com/metacoglab/meta_dots). On each trial of this task, two circles enclosing sets of dots appeared for 200 ms on either side of a central fixation cross (each circle with a radius of 5 degrees of visual angle, located along the middle of the screen, with an eccentricity from the vertical midline of 5.5 degrees of visual angle). One of the two circles always contained 50 dots while the other varied in dot number, and the position (left/right) of the circles was randomized on each trial. In a 2-alternative forced-choice (2AFC) task, participants discriminated which of the circles contained more dots by pressing the left or right arrow keys on the keyboard. The difference in the number of dots was determined by a pair of interleaved 2-down-1-up adaptive staircases aimed at fixing performance at around 71% accuracy. After each response, participants reported their confidence in the accuracy of their own response using the same vertical visual analog scale that they had used for the two previous tasks rating confidence and similarity for facial expressions.

Before the main visual task, we ran 80 trials of a staircase procedure where participants did only the discrimination task without rating confidence. Here we also included two interleaved 2-down-1-up staircases starting from a difference of 3 and 20 dots respectively. One participant (unintentionally) received feedback about the accuracy of the discrimination task while rating confidence, so we excluded their data from the analysis. The visual task took approximately 20 minutes. Over all participants, we also excluded 2% of the trials where the reaction times to either the discrimination task or the confidence rating were faster than 300 ms or slower than 5 s. We estimated metacognitive efficiency as Mratio^30^ after scaling and binning confidence into four discrete confidence levels based on uniform intervals.

### Toronto Alexithymia Scale

At the end of the experiment we collected responses to a computerized version of the Toronto Alexithymia Scale (TAS^62^) running on a browser, and the data were stored locally^63^ (jatos.org). Most participants completed a German version of the scale, except for seven non-German speakers who completed an English version instead. The TAS-20 consists of 20 items that can each be answered on a 5-point Likert scale. We considered three out of the four subscales (Difficulty identifying feelings, Difficulty describing feelings, and Externally-oriented thinking, but excluded the Daydreaming subscale). We calculated Bayes Factors (BF_10_) for correlations between these covariates and individual slopes from the estimated models using the *BayesFactor* package^64^ in R (version 3.6.2).

### Data processing and analysis

Following the pre-registered plan, we excluded trials from the facial expressions task at the single participant level if RTs (time between image onset and key press) were above the 95 percentile for that participant. This cutoff was necessary because we noticed that participants sometimes laughed at their own picture or got otherwise distracted. This resulted in seven trials excluded from the entire dataset where the time to take a picture was below 300 ms, and a mean lower threshold of exclusion of 9.43 s (range: 4.0 - 18.0 s).

For each of the pictures taken, we obtained the x,y coordinates of landmarks distributed on the face. In our pre-registered plan we stated that we would estimate the landmark positions using two different toolboxes and choose the best one to estimate distance based on the quality of the relationship to the similarity ratings. Instead, due to technical problems in running one of the toolboxes we opted for the Face Alignment package^65^ alone (https://github.com/1adrianb/face-alignment v.1.0.0), a fully automated deep-learning based face alignment network (FAN) that places landmarks on the pictures. We used the *face-alignment* package together with *scikit-image and pytorch* to extract the landmarks from the faces, running on Python v3 in a Jupyter notebook v5. The face-alignment package automatically places 68 landmarks on the face and excludes the forehead and hairline.

Using MATLAB (R2020a), we computed the distance (in coordinate space) between each pair of target and response images. Using the x,y coordinates for all landmarks, we ran a Procrustes rigid alignment of each face in a pair to a standardized set of coordinates. We used three minimally variant reference points for this alignment: the outer corners of each eye and a point just below the nose. The transformation allowed for translation, orthogonal rotation, and scaling. Thus, these linear transformations minimized the variance in the distance data that could be accounted for by head rotations and general enlargement or shrinkage due to change in the face position. It did not account for other rotations (yaw and pitch), where the relative distance between some face components can change without the facial expression being different. After rigid transformation, we calculated the total distance for each pair of target and response images as the Euclidean distance (the root of the sum of squares, see equation in Figure 1) over all 68 landmarks between the two images. We refer to this measure simply as the distance between two images. We then log-transformed the obtained distances to ensure that the data were normally distributed before fitting the Bayesian mixed models.

### Bayesian mixed models

In our central analysis we computed metacognitive access to facial expressions as the relationship between confidence ratings and performance. We take the distance as an inverse measure of performance: if a response image closely matches the target image, the distance between them will be small. Furthermore, a strong negative relationship between confidence ratings and distance will indicate that participants had metacognitive access to their own facial expressions, as they (correctly) provided low ratings in trials where the two images differed the most. Conversely, no relationship between confidence and distance would indicate that participants had no metacognitive access to their own expressions.

Because finding no relationships between variables was a plausible outcome from our analyses, we used Bayesian statistics that, unlike frequentist statistics, provide evidence for the null hypotheses. We analyzed the data using Bayesian mixed models created in Stan (http://mc-stan.org/) through the *brms* package^66, 67^. In all cases, we ran 4 chains with 15,000 iterations, 5,000 burn-in samples each, and no thinning. We checked for convergence by visually examining the MCMC chains and ensured that the scale reduction factor (Rhat) of all models was equal or close to 1. We considered that ratings might vary across participants both in their mean and in their relationship to the landmark distance, and that different facial expressions might vary in their associated difficulty to both reproduce (leading to greater variability in the landmark distance) and to rate (leading to differences in the ratings). Thus, in all models and unless otherwise stated, we included random slopes for both participants and facial expressions (see the explicit model syntax in Table 1). We extracted the participant-wise random slopes using the *mixedup* package (https://m-clark.github.io/mixedup/).

**Table 1:**
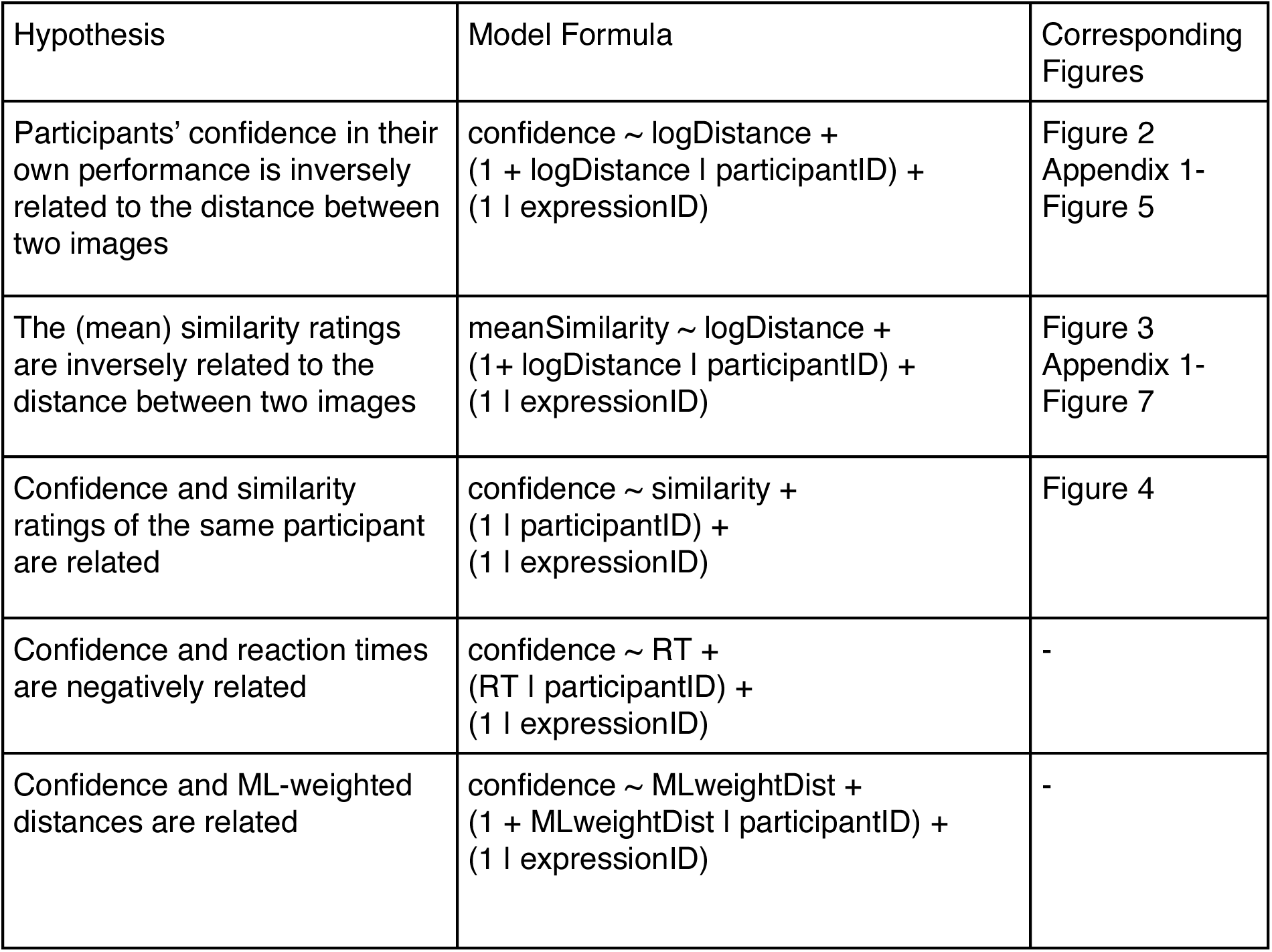
Formulas for the Bayesian mixed models employed

Because, to the best of our knowledge, there was no existing data to inform our priors, we followed recommendations^68^ to use heuristics to define prior distributions. We built the prior for the slope between ratings and distance based on the ratio-of-scales heuristic: we found that the range of (log-transformed) distances was approximately 3 a.u. (arbitrary units), whereas the range of confidence ratings is 1 point (minimum: 0). Therefore we used a normal prior centered on 0 with an SD = ⅓ (which corresponds to the ratio between confidence range and distance range) for the slope parameter. To find a prior for the model intercept we followed the logic behind the room-to-move heuristic. Note that raw distances ranged between [131.36 - 2493.78] a.u., hence the expected rating at 0 distance (i.e., perfect performance) can be well approximated by the expected rating at distance = 1, which corresponds to the intercept in a linear model with log-transformed distances. We reasoned that a participant with maximum metacognitive performance would consistently rate their confidence as 1, when the distance between the two images was 0. Because we realistically expect participants to have (at most) less than perfect metacognitive access to their own expressions, we centered the prior at 0.8 with an SD = 0.5. Following a similar logic, we set the prior slope between the two ratings to be centered at 0 with SD = 1, and an intercept of 0 with an SD = ½. For all models, we report the estimate, its associated error mean, the 95% credibility interval (CI), and the BF_10_, estimated using the *bayestestR* package^69^, to compare each model against its null counterpart, containing the same random effects structure but not the fixed effect of interest. We also include the posterior draws for each participant in relation to the region of practical equivalence (ROPE). We set the ROPE to a default range from −0.1 to 0.1 of a standardized parameter, which corresponds to a negligible effect size^70, 71^. Finally, we estimated R^2^ values as implemented by the *brms* package^72^.

We computed metacognitive access to faces using only linear regression and estimated the correlation with visual Mratios, deviating from the pre-registered plan. We initially planned to also calculate the area under a type-2 ROC curve (AUROC2) by arbitrarily assuming that first-order performance on the Faces task was at 70% accuracy and by classifying trials with distances above the corresponding threshold as “incorrect”. This analysis had the advantage that it would have allowed us to correlate metacognitive performance measured on the same scale for both tasks (Faces and Visual), but we later reasoned that it would make the results less easily interpretable while not adding explanatory power and therefore decided to omit it.

### Machine learning models

Using Python v3, and *scikit-learn*, we created a separate model for each subject wherein, first, each landmark distance was determined by (x,y) coordinate differences between the two images. We further decomposed the differences into four zero- or positive features (one for each cardinal direction). This allowed different directions of movement to be weighted differently by the model. We normalized each feature by dividing it by its median. Then, we applied dimensionality reduction using principal component analysis with a set number of principal components (66, or approximately 90% of the variance from all subjects) in order to avoid multicollinearity among the features. Finally, a least squares linear regression model was trained for each participant using trial-wise leave-one-out cross-validation.

The resulting ML model weights referred to features in principal component space. We translated the model weights back into landmark space (i.e., x,y coordinates of the facial landmarks). To do so, we approximated the weight *w* of each feature *f* using the expression in (1):

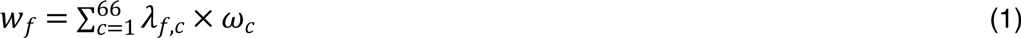

Where *λ_*f*,*c*_* is the loading of feature *f* on principal component *C*, and *ω*_*c*_ is the ML model’s weighting of principal component *c*.

To reconstruct the distances weighted by the results of each ML model, we used expression (2):

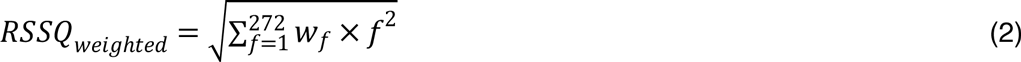

Where *w_f_* denotes the weights for each feature *f*, which is in turn the difference between response and target images for each cardinal direction, for a given landmark, if the difference was positive, and 0 otherwise. The 272 features result from decomposing 68 landmarks into the four cardinal directions. Note that unlike the case for the Euclidean distance, where distances were forced to be positive and each of them had an effective weight of 1, here we allowed the feature weights to be signed. For those cases where the term under the square root was negative, we calculated the root of the absolute value and then reversed the sign. Note that *RSSQ_weighted_* is now better interpreted as a measure of performance, and not distance: because the ML-derived weights already account for the negative relationship between distance and confidence, *RSSQ_weighted_* is expected to show a positive relationship to confidence.

We obtained adjusted R2 for each (participant-specific) model values and compared them using a Bayesian Wilcoxon Signed-Rank test^73^ as implemented in JASP^74^ v0.14 with 10,000 MCMC samples and 5 chains, and a default Cauchy prior.

## Acknowledgements

We thank student assistants for help in data collection in Experiment 1, and Manuel Zellhöfer for help in programming the experimental paradigm. We thank Soledad Galli for assistance with the ML models and Nathan Faivre for comments on an earlier version of this manuscript. ABC, CF and EF were supported by a Freigeist Fellowship to EF from the Volkswagen Foundation (grant number 91620). This work was supported by the Deutsche Forschungsgemeinschaft (DFG, German Research Foundation) - 337619223 / RTG2386 and the Max-Planck Society. The funders had no role in the conceptualization, design, data collection, analysis, decision to publish, or preparation of the manuscript.

## Data and Code Availability

Raw data (excluding images from participants and any other personally identifiable information) along with reproducible analysis scripts are available under https://gitlab.com/elisa.filevich/cistonetal_metacognitionoffacialexpressions.

## Appendix 1

### Supplementary Information

Appendix 1-Figures 1-3 show the single-trial data (and linear regressions at the single-participant level) for the data reported in the main text.

**Appendix 1-Figure 1:**
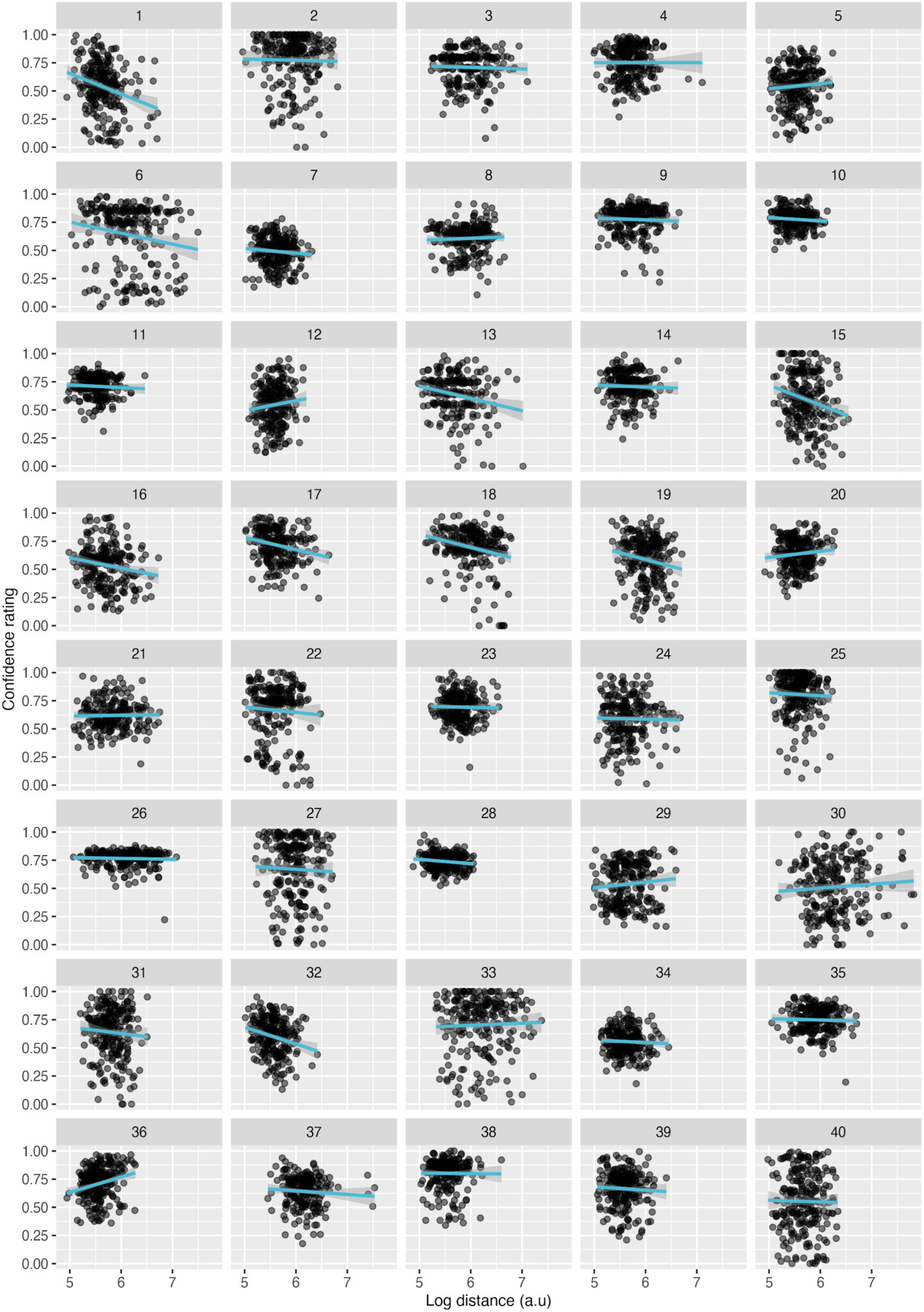
Linear regressions of confidence ratings as a function of distance. (at the single-participant level, for Experiment 2). Note that all statistical inferences are made on the basis of Bayesian linear regressions, and this plot is for illustrative purposes only.

**Appendix 1-Figure 2:**
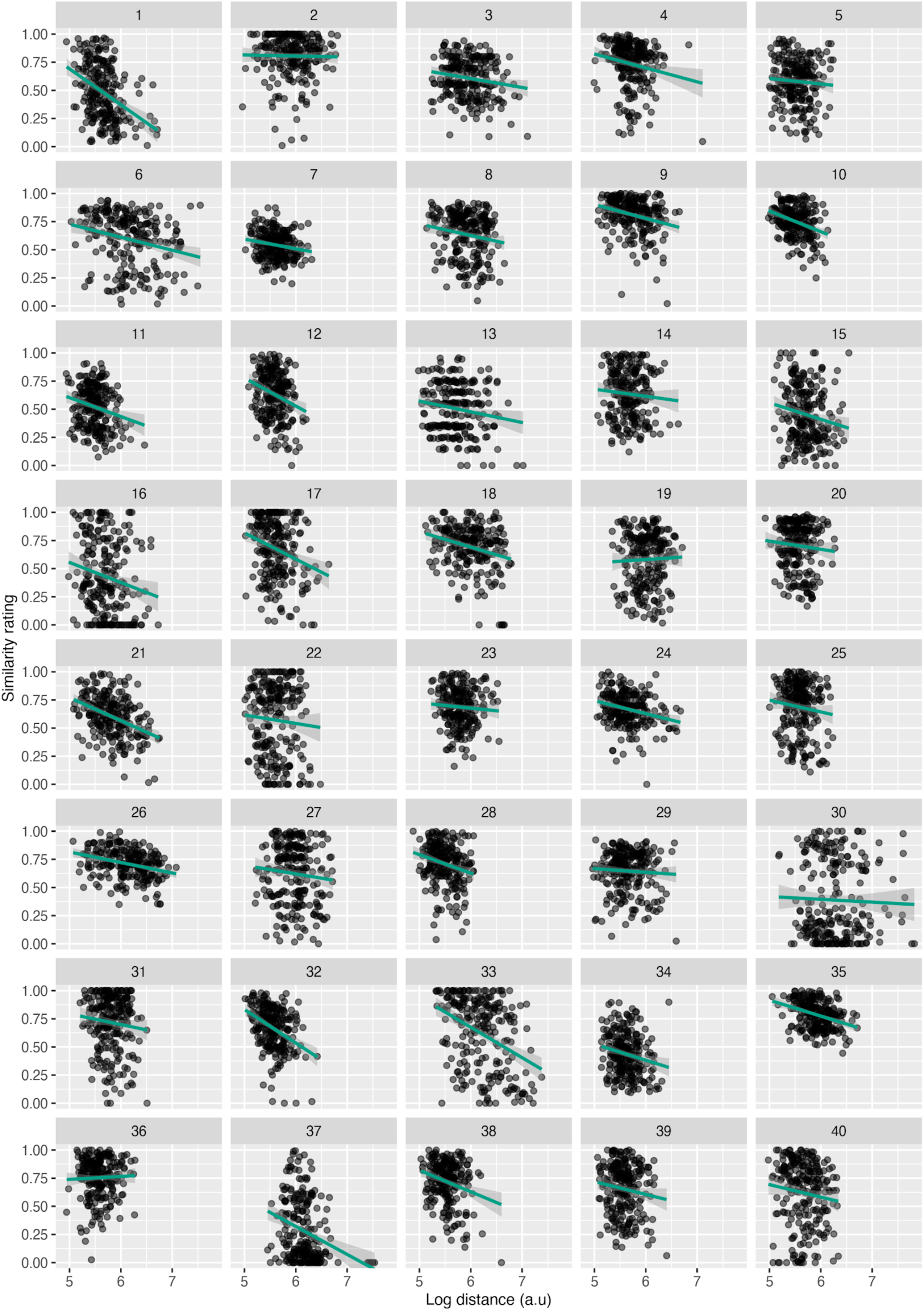
Linear regressions of similarity ratings as a function of distance. (at the single-participant level, for Experiment 2). Note that all statistical inferences are made on the basis of Bayesian linear regressions, and this plot is for illustrative purposes only.

**Appendix 1-Figure 3:**
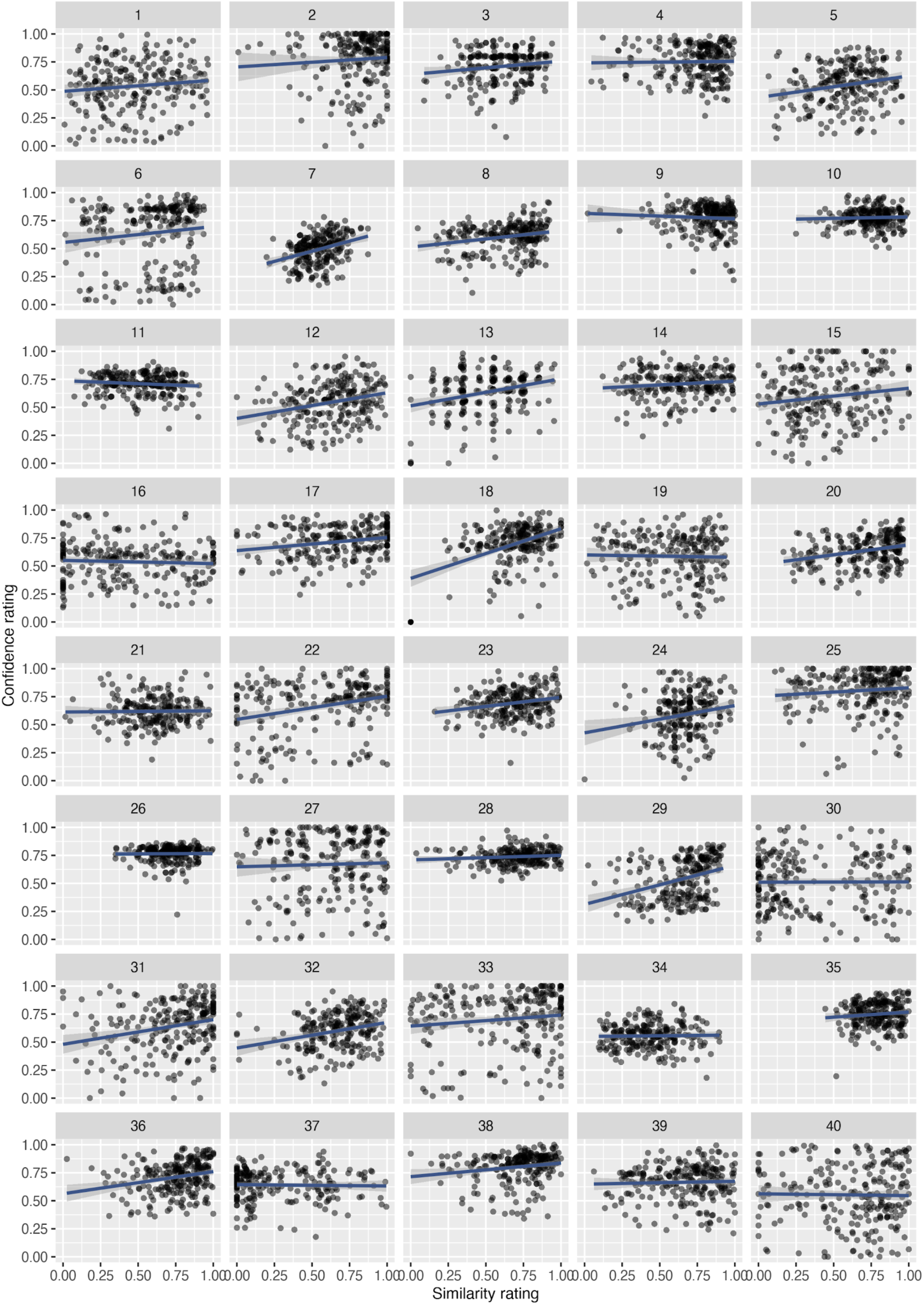
Linear regressions of confidence vs. similarity ratings. (at the single-participant level, for Experiment 2). Note that all statistical inferences are made on the basis of Bayesian linear regressions, and this plot is for illustrative purposes only.

### Supplementary analyses

#### Machine Learning - Effects of RT

Because confidence is known to correlate negatively with response times^75, 76^ (RT), we first explored a potential relationship between the two and asked whether RTs could have served as a proxy for performance. We ran a Bayesian linear regression model of participants’ confidence ratings including the RT as a fixed effect and random intercepts for participant and facial expression, as well as a per-participant random slope for RT. We based the prior distribution for this analysis on previous data from our group^3^, and set a wide prior for the intercept centered at around confidence = 0.8 with SD = 0.5, and a prior for the slope centered on 0 with an SD = 0.20, which roughly corresponds to the ratio-of-scales. We confirmed that there was a small but consistent effect of RT on confidence (M = −0.01 ± 0.00, CI = [−0.02, −0.00], BF_10_ = 5.66 x 10^39^, R^2^ = 0.20).

To evaluate whether the landmarks informed confidence ratings above and beyond RT, we compared the resulting individual r values from the ML models (including both RTs and the x, y positions of the landmarks) to those of a ML model including only RTs as their single feature (Appendix 1-Figure 4). A non-parametric ANOVA computed with the ez package for R revealed an interaction effect (*p* = 0.001) on the *r* values between the variable predicted (confidence or similarity) and the features included in the model. Note that we do not interpret the main effect of the number of features included, as these are known to inflate the r values. Instead, we focus on the interaction effect. In particular, the interaction revealed higher *r* values for the models of similarity that included both landmarks and RT as compared to confidence (Wilcoxon signed rank test, *p* = 0.015), but lower *r* values for models including RTs only (Wilcoxon signed rank test, *p* = 0.068). This pattern of results is consistent with landmarks being predictive of confidence ratings, above and beyond RTs To understand the contribution of RTs relative to the other features, we obtained the rank of importance of RTs within the ML model. We found that RTs varied in importance with each participant, but ranged between the 5th and the 100th percentile (Mean = 71.15, Median = 93.41), suggesting that, for some participants, RTs were the most reliable piece of information for confidence ratings, even if the variance explained by them was very low.

**Appendix 1-Figure 4:**
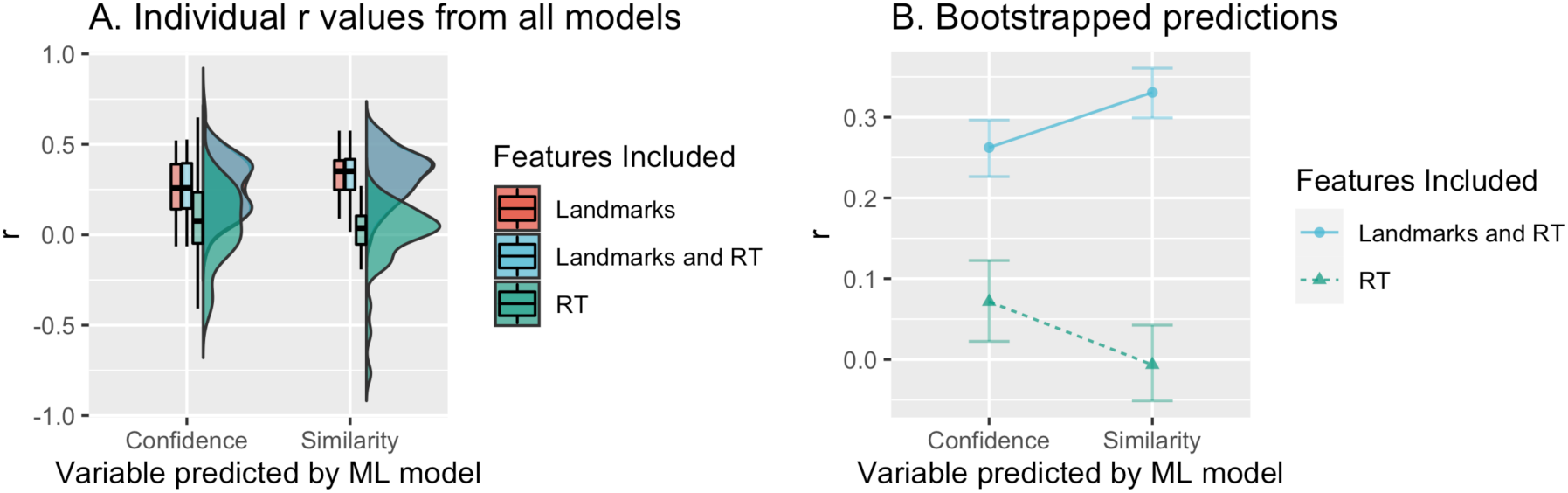
r values resulting from the linear regression models built using ML. **A.** Distributions of individual r values (summarized with boxplots and violin plots) for models on confidence or similarity ratings, using different sets of features. **B. Bootstrapped predictions from a non-parametric ANOVA** for models of confidence and similarity built using RTs alone or also including landmark information.

### Supplementary Pilot Experiment

Prior to pre-registering and collecting the data reported in the main text, we collected a smaller dataset as a pilot. Because there are some important differences in the experimental details, we report the methods and results as supporting information, that serve as a conceptual replication.

### Supplementary Methods - Pilot Experiment

The methods for the pilot experiment were largely similar to those of the main experiment. We only describe here the differences between the two.

#### Participants

Thirteen healthy participants took part in the experiment after giving informed consent (seven female, mean ± SD: 24 ± 3 years). One participant was excluded from the analysis because four external judges agreed (see below) that there was no variability in their facial expressions. Participants had no recent history of psychiatric disease. The local ethics committee approved all procedures, which conformed to the Declaration of Helsinki.

#### Apparatus

Behind the mirror, a digital camera (Logitech HD C310) connected to the computer captured images of the participants’ facial expressions. The apparatus was similar to the one described in the main text, with some minor differences. Unlike in the main experiment, where the screen rested on top of the stimulus box and projected downwards, the screen lay on the table for the pilot experiment and projected upwards. From the point of view of the participants, this did not change the visual display.

#### Procedure

The task was programmed on GNU Octave and displayed stimuli using Psychtoolbox-3^58–60^, and ran on a Linux Debian (Gnome 3.4.2) operating system. The task consisted of two parts (not three). Participants saw 30 (not 32) different photos of four different actors and imitated each expression as best they could. The images were presented in one of five possible pre-defined random orders to each participant. As in the main experiment, participants first generated 30 participant-specific pictures that then served as target images for the second part of the paradigm. After each trial, participants rated their confidence (on a scale from 1 to 6) regarding how well they thought that they had imitated their own previous expression. To make the task intuitive, we kept the mapping of the scale consistent with the German education system, where the best grade is a 1.0. We then reversed the ratings for further analyses, so that a rating of 6 corresponds to the highest confidence. In all cases, we recorded each picture taken, the response time (RT, measured as the time between image onset and key press) and participants’ confidence ratings. Participants saw each of their 30 target expressions repeated 8 times in random order, for a total of 240 trials. We only revealed that they would have to reproduce their own expressions after the first part of the experiment was complete. On average, the experiment took approximately 50 minutes.

#### Data Processing and Analysis

We first used the Face Modeling GUI^78^ to manually position 99 landmarks on their corresponding locations on a small subset of images (3-5) of each participant. The Face Modeling GUI then uses the location of these landmarks to automatically find their optimal locations in the remaining images. After the automatic fit, the landmarks in each of the images were corrected manually. In this way, we reduced the dimensionality of each of the 240 response images along with the 30 target images for each of the participants to 99 pairs of (x,y) coordinates. We then did the same Procrustes rigid-alignment as described in the main text, with 5 reference points instead of 3 (the inner and outer corners of each eye and a point just below the nose). We did not use a mean reference face, but instead minimized the distance of each response picture to its corresponding target picture.

#### Similarity ratings by external judges

Unlike what was the case in the main experiment, here four independent judges (student research assistants) rated the image pairs for similarity on a scale from 1 to 6, exactly like the one the participants had used.

#### Data processing and analysis

Here as well we followed recommendations^68^ to use heuristics to define prior distributions. We built the prior for the slope based on the ratio-of-scales heuristic: we found that the range of (log-transformed) distances was approximately 4.93 a.u. (arbitrary units), whereas the maximum possible range of ratings is 5 points (maximum: 6, minimum: 1). The ratio between the two is approximately 1, so we used a normal prior centered on 0 with an SD = 1 for the slope parameter. To find a prior for the model intercept (the expected rating at 0 distance, i.e., perfect performance), we followed the logic behind the room-to-move heuristic. We reasoned that a participant with maximum metacognitive performance would consistently rate their confidence as 6, when the distance between the two images was 0. Because we realistically expect participants to have (at most) less than perfect metacognitive access to their own expressions, we centered the prior at 4 with an SD = 3.

### Supplementary Results - Pilot Experiment

Because ratings were not on a visual analog scale but instead on a Likert scale, we first quantified our participants’ metacognitive access to their own facial expressions using an ordinal Bayesian mixed-effects regression model of participants’ confidence ratings. The model included the log-transformed landmark distances as a fixed effect (for all 99 landmarks combined) as well as random intercepts for participant and facial expression (See Appendix 1-Table 1). The estimated ordinal regression coefficient was indistinguishable from 0 (M = 0.04 ± 0.07, CI = [−0.10, 0.16]) and the evidence ratio favoured the (point) null hypothesis of no relationship between confidence and distance (BF_10_ = 0.082). This is illustrated by the flat probability profiles for each rating shown in Appendix 1-Figure 5.A: while there were differences in the overall probability of each confidence rating (e.g. a rating of 5 occurring more often than others), the probability of a participant providing a given confidence rating was similar over all landmarks distances (see also Appendix 1-Figure 6 for the single-participant data).

**Appendix 1-Figure 5.**
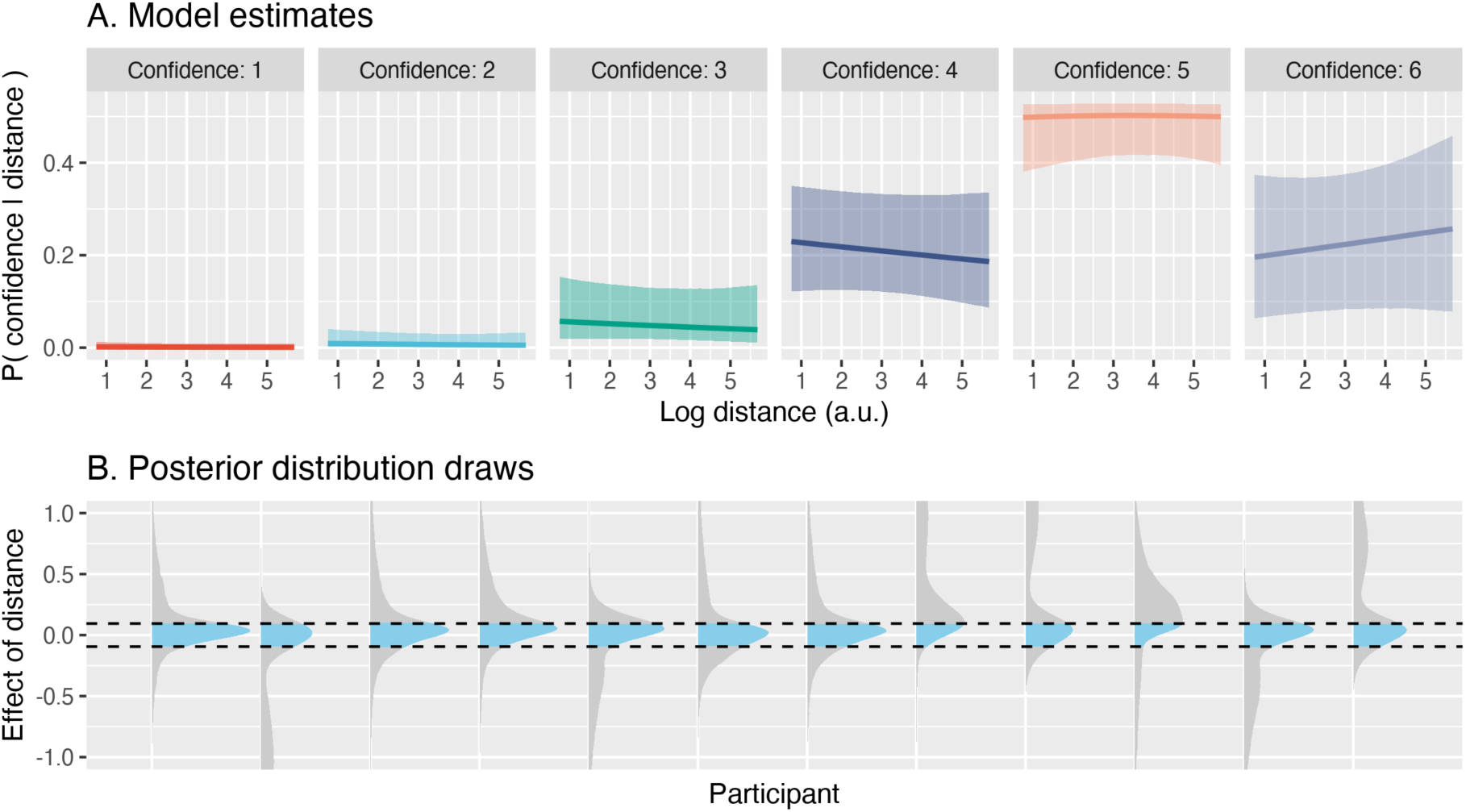
No evidence for metacognitive access to facial expressions. **(A.)** Group effects reflecting mean metacognitive access, namely the relationship between confidence ratings and distance between two images (inverse of performance). While different confidence ratings appear at different frequencies in the data, they do not vary with distance as would be expected if participants had metacognitive access to their own expressions. Solid lines represent the mean of the posterior draws, the shaded regions represent the 95% credibility interval. **(B.)** Posterior draws for each subject, shown in relation to the ROPE. Note that the y-axis is clipped to better display the distributions around the ROPE and therefore excludes the long tails of some of the distributions.

**Appendix 1-Figure 6:**
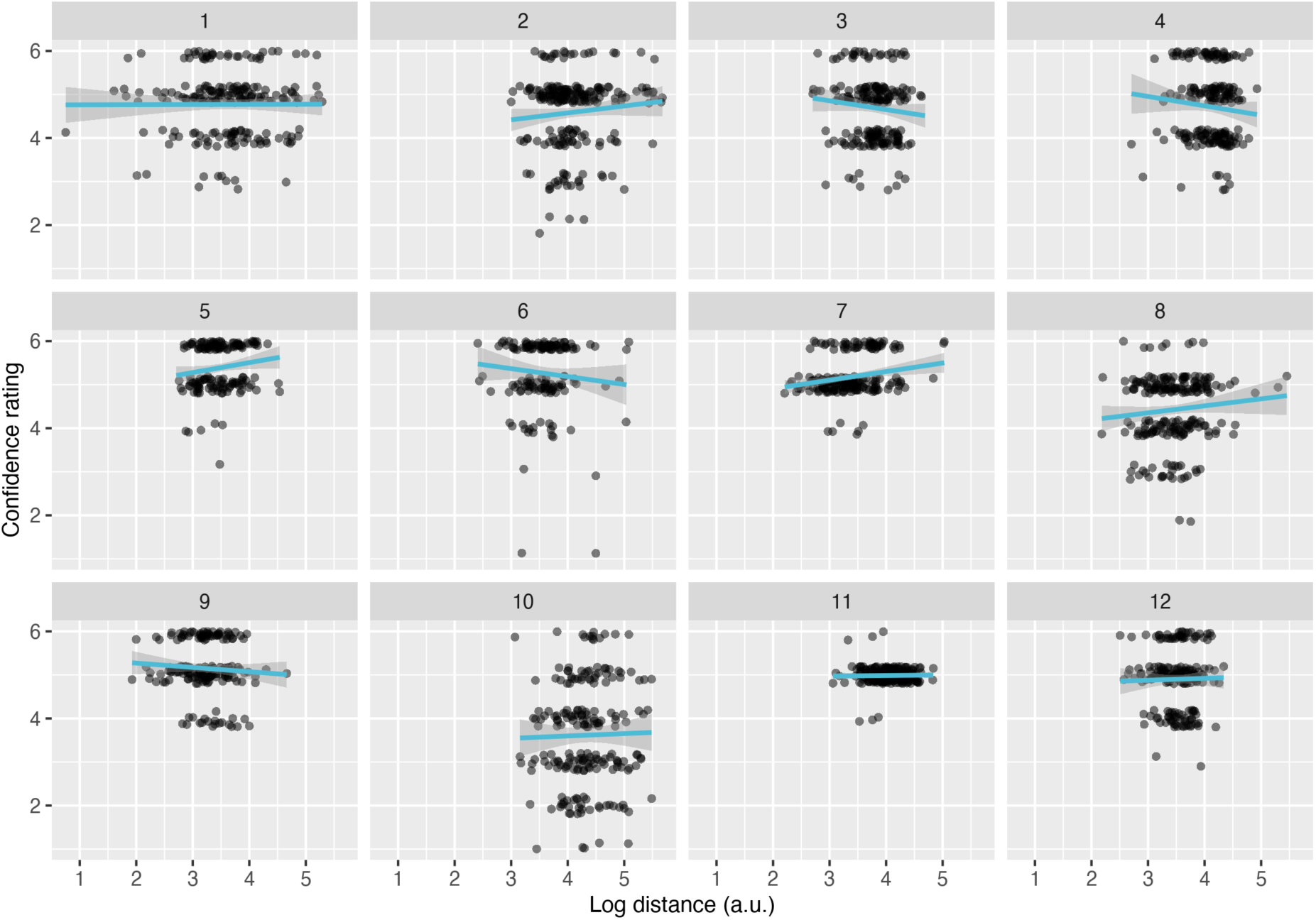
Linear regressions of confidence ratings as a function of distance. (at the single-participant level, for Experiment 1). Note that all statistical inferences are made on the basis of Bayesian ordinal regressions, and this plot is for illustrative purposes only.

**Appendix 1-Table 1:** Bayesian ordinal model estimates for the effect of distance on similarity. Each row contains the estimates for a single judge (and all participants in the pilot experiment) and includes the mean, standard deviation, 95% credibility interval and BF_10_ relative to the null model.

| Judge | Effect of distance on similarity rating ($M \pm SD$) | 95% CI | $BF_{10}$ |
| --- | --- | --- | --- |
| 1 | -0.53 ± 0.11 | [-0.75, -0.32] | 554.91 |
| 2 | -0.55 ± 0.07 | [-0.70, -0.41] | 15175.95 |
| 3 | -0.89 ± 0.11 | [-1.11, -0.67] | 57346.06 |
| 4 | -0.73 ± 0.11 | [-0.95, -0.51] | 7608.01 |

That there is no observable relationship between the combined landmark distances and participants’ confidence ratings suggests, at face value, that participants did not have access to the details of their face. However, other alternative explanations must be considered. First, it is possible that the landmark distance measure, which is essentially the result of an algorithm placing landmarks based on pixel information plus some rigid transformations, may not capture enough information relevant for the similarity of two faces. If this were true, there should also be no relationship between the landmark distance and the similarity ratings provided by external judges looking at each image pair side by side. In fact, this was not the case. To evaluate this possibility we used a Bayesian linear mixed-effects regression model on the mean of four judges. The model included the same fixed and random effects factors as in the mixed ordinal model above (namely, the log-transformed distance as a fixed effect, intercepts for participant and expression as random effects, and a by-participant random slope for the fixed effect). However, unlike in the mixed-effects regression model on participants’ confidence ratings, we did find a consistent negative relationship between the distance and the similarity ratings (M = −0.54 ± 0.06, CI = [−0.67, −0.42], BF_10_= 71551.85). That is, unlike the confidence ratings, the similarity ratings did show a consistent and (as expected) negative relationship to the distance (Appendix 1-Figure 7.B and Appendix 1-Figure 8). This suggests that the distance did carry some information about face similarity meaningful to human observers. For illustration purposes only, we repeated the analysis between similarity ratings and distance but this time rounded the mean ratings and ran an ordinal model (Appendix 1-Figure 7.B). We do not make any statistical inferences from this analysis but use it only to illustrate the differences between the probability profiles of the ratings that vary with distance and those who do not (Appendix 1-Figure 5.A).

**Appendix 1-Figure 7.**
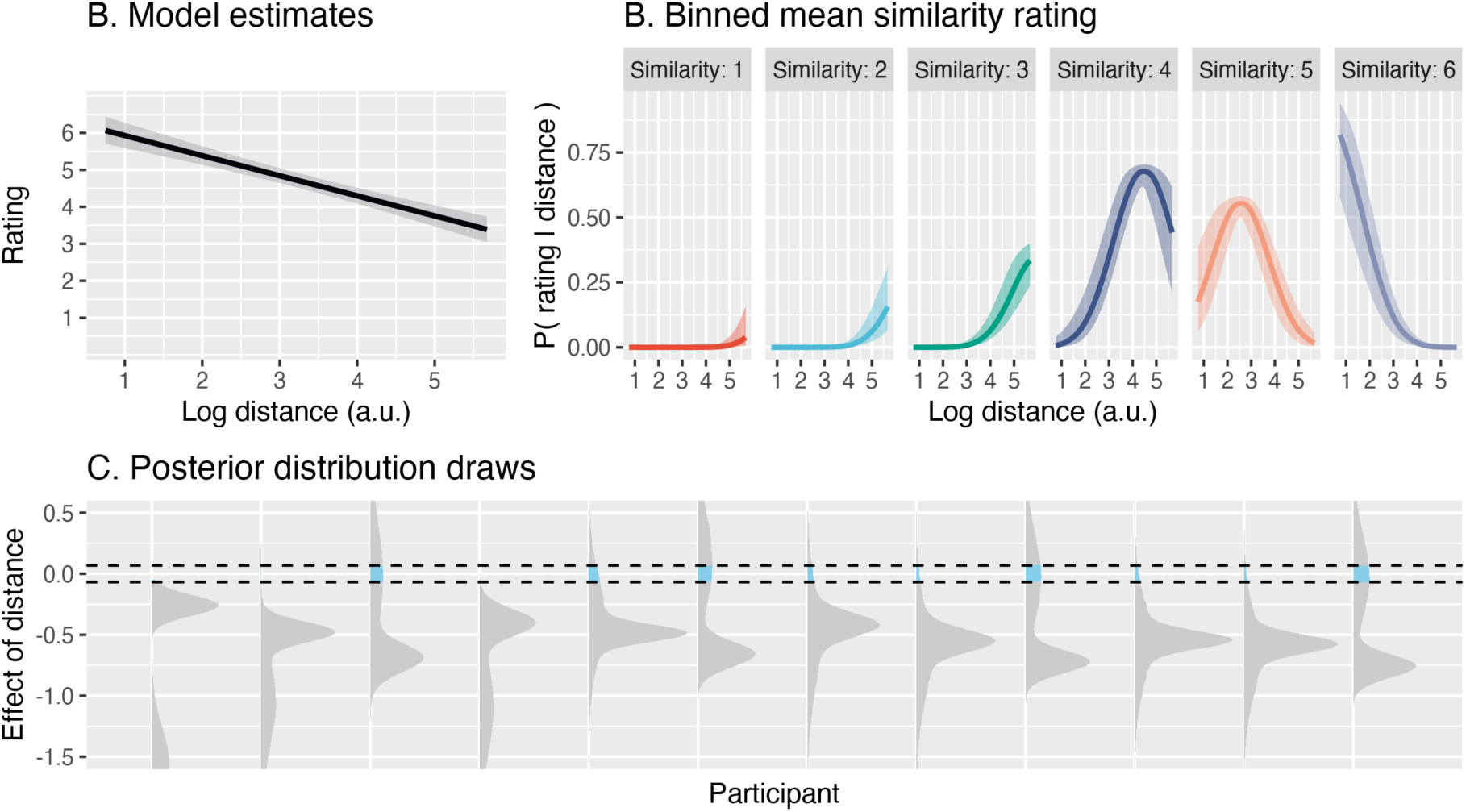
The distance between two images captures relevant information. **(A.)** Group effects reflecting the information contained in the distance between two images, namely the relationship between the mean similarity ratings of four judges (who viewed each image pair side-by-side) and distance between two images. There is a clear relationship between mean similarity and distance, suggesting that distance contains meaningful variability. **(B.)** An ordinal version of the model shown in (A.) presented only to illustrate the contrast to Appendix 1-Figure 5. For both panels (A.) and (B.), solid lines represent the mean of the posterior draws, and the shaded regions represent the 95% credibility interval. **(C.)** Posterior draws for each subject, shown in relationship to the region of practical equivalence (ROPE). Note that the y-axis is clipped to better display the distributions around the ROPE and therefore excludes the long tails of some of the distributions.

**Appendix 1-Figure 8:**
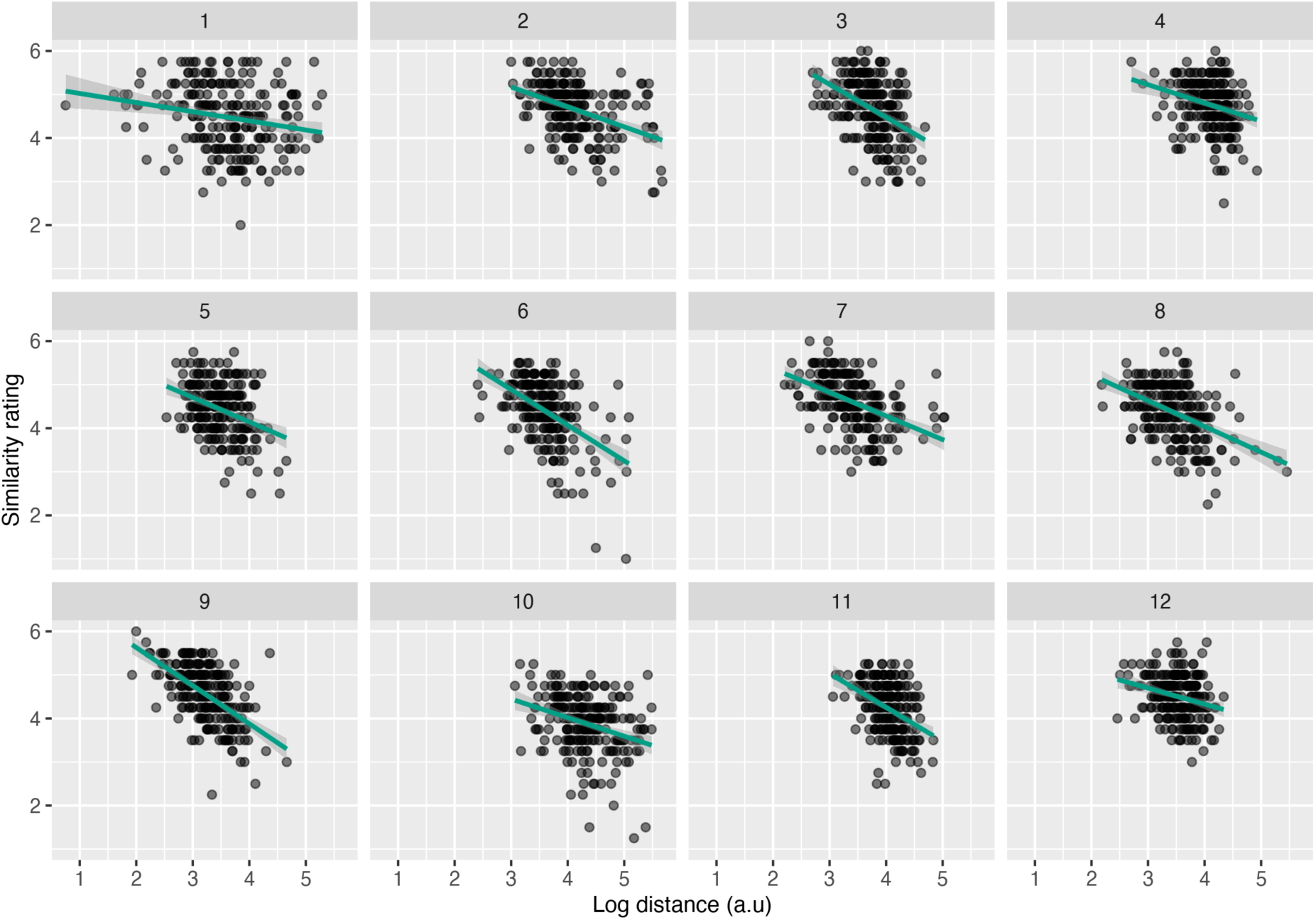
Linear regressions of mean similarity ratings as a function of distance. The y axis represents the mean of all four judges, and each panel represents a single participant, from the pilot experiment).

As in the main experiment, here we also found that distance was related to similarity ratings. Neither the procedure to estimate distance nor the similarity ratings were identical between the two experiments (two different algorithms placed 68 or 99 landmarks respectively; and either the participants themselves or external judges rated similarity), which validated our measure of distance by showing that it does not depend on idiosyncratic properties of the algorithm or the rating process.

Importantly, we note that the relationships shown in Appendix 1-Figure 7, panels B. and C. and Appendix 1-Figure 8 are the result of taking the mean of four judges. Thus, this significant relationship might be accounted for by a Wisdom of the crowds effect, whereby the mean of the estimates of many individuals is better than any single individual’s estimate^79^. To evaluate this possibility, we ran Bayesian ordinal mixed regressions for the similarity ratings of each individual judge. In all cases, we found that the estimates were negative, and clearly different from 0 (all mean slope estimates < −0.53, all BF_10_ > 554. See Appendix 1-Table 1 and Appendix 1-Figure 9 for the model predictions and single-participant data, respectively).

**Appendix 1-Figure 9:**
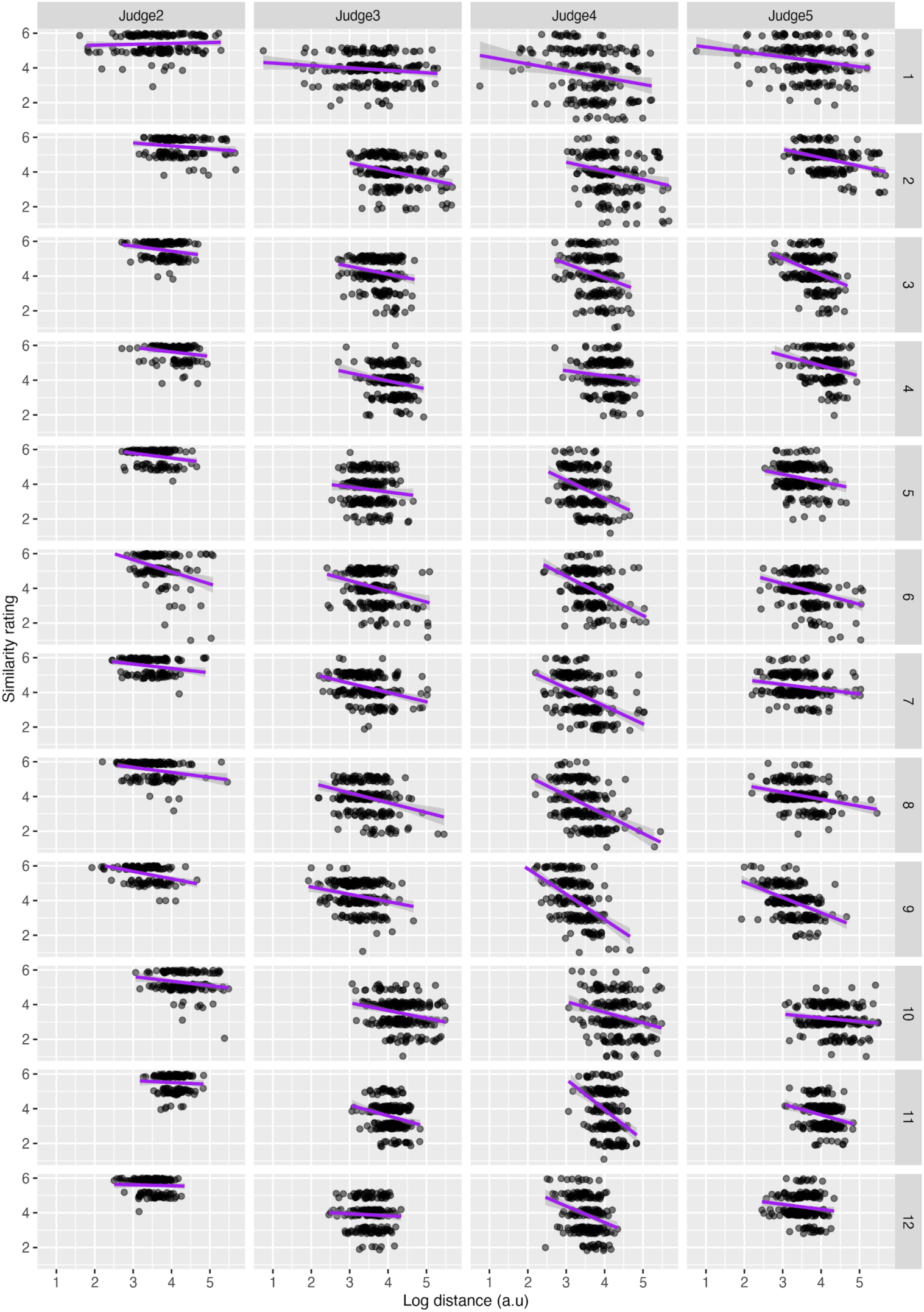
Linear regressions of similarity ratings from each judge as a function of distance. Each panel represents a single judge (columns) and participant (rows) from the pilot experiment.

### Brief Discussion - Pilot Experiment

Briefly, these results suggest that participants did not have access to the low-level details of their own facial expressions. This could not be explained by any of the several alternatives we explored: neither lack of variability in performance or a poor benchmark measure (similarity ratings from external judges did show a clear relationship to the landmark distances, Appendix 1-Figure 7) nor the fact that the confidence ratings were from a single person (individual judges’ similarity ratings also showed the same clear relationship, Appendix 1-Table 1) proved to be sufficient to explain the apparent null relationship between confidence ratings and distance.

Despite these controls, alternative explanations remain in principle possible, which we incorporated when designing the experiment reported in the main text. First, participants provided their confidence ratings on a Likert scale from 1-6. Perhaps, a continuous scale would have given them the opportunity to provide more nuanced and precise ratings. Second, metacognitive ability — in both the visual^80^ and the motor^41^ domains — is known to vary in the normal population. Perhaps, due to mere chance, participants with poor general metacognitive access to their own facial expressions were overrepresented in the relatively small sample of 12 participants. Hence, to exclude the possibility that our conclusions in this pilot experiment resulted from a small (and potentially biased) sample of 12 participants, we tested a larger sample. Third, we considered the possibility that the differences we observed in this pilot experiment between the relationships of distance and confidence and similarity ratings could be attributed to differences in metacognitive traits between groups of individuals. We therefore did not recruit external judges but asked the same participants to rate their own performance in the image pairs.

## References

1. Kal, E., Prosée, R., Winters, M. & Kamp, J. van der. Does implicit motor learning lead to greater automatization of motor skills compared to explicit motor learning? A systematic review. PLOS ONE 13, e0203591 (2018).

2. Kleynen, M. et al. Using a Delphi technique to seek consensus regarding definitions, descriptions and classification of terms related to implicit and explicit forms of motor learning. PloS One 9, e100227 (2014).

3. Taylor, J. & Ivry, R. Implicit and Explicit Processes in Motor Learning. 63–87 (2013) doi:10.7551/mitpress/9780262018555.003.0003.

4. MacIntyre, T., Igou, E. R., Campbell, M. J., Moran, A. P. & Matthews, J. Metacognition and action: a new pathway to understanding social and cognitive aspects of expertise in sport. Front. Psychol. 5, (2014).

5. Proske, U. & Gandevia, S. C. The Proprioceptive Senses: Their Roles in Signaling Body Shape, Body Position and Movement, and Muscle Force. Physiol. Rev. 92, 1651–1697 (2012).

6. Sherrington, C. S. The integrative action of the nervous system. (Scribner, 1906).

7. Tuthill, J. C. & Azim, E. Proprioception. Curr. Biol. 28, R194–R203 (2018).

8. Goodwin, G. M., McCloskey, D. I. & Matthews, P. B. The contribution of muscle afferents to kinaesthesia shown by vibration induced illusions of movement and by the effects of paralysing joint afferents. Brain J. Neurol. 95, 705–748 (1972).

9. Lackner, J. R. SOME PROPRIOCEPTIVE INFLUENCES ON THE PERCEPTUAL REPRESENTATION OF BODY SHAPE AND ORIENTATION. Brain 111, 281–297 (1988).

10. Craske, B. & Crawshaw, M. Shifts in kinesthesis through time and after active and passive movement. Percept. Mot. Skills 40, 755–761 (1975).

11. Fuentes, C. T. & Bastian, A. J. Where is your arm? Variations in proprioception across space and tasks. J. Neurophysiol. 103, 164–171 (2010).

12. Gritsenko, V., Krouchev, N. I. & Kalaska, J. F. Afferent input, efference copy, signal noise, and biases in perception of joint angle during active versus passive elbow movements. J. Neurophysiol. 98, 1140–1154 (2007).

13. Limanowski, J. & Blankenburg, F. Integration of Visual and Proprioceptive Limb Position Information in Human Posterior Parietal, Premotor, and Extrastriate Cortex. J. Neurosci. 36, 2582–2589 (2016).

14. Ruttle, J. E., Hart, B. M. ‘t & Henriques, D. Y. P. The fast contribution of visual-proprioceptive discrepancy to reach aftereffects and proprioceptive recalibration. PLOS ONE 13, e0200621 (2018).

15. Sober, S. J. & Sabes, P. N. Flexible strategies for sensory integration during motor planning. Nat. Neurosci. 8, 490–497 (2005).

16. van Beers, R. J., Wolpert, D. M. & Haggard, P. When Feeling Is More Important Than Seeing in Sensorimotor Adaptation. Curr. Biol. 12, 834–837 (2002).

17. van Beers, R. J., Sittig, A. C. & van der Gon Denier, J. J. How humans combine simultaneous proprioceptive and visual position information. Exp. Brain Res. 111, 253–261 (1996).

18. Clark Weeden, J., Trotman, C.-A. & Faraway, J. J.. Three Dimensional Analysis of Facial Movement in Normal Adults: Influence of Sex and Facial Shape. Angle Orthod. 71, 132–140 (2001).

19. Coulson, S. E., Croxson, G. R. & Gilleard, W. L. Quantification of the Three-Dimensional Displacement of Normal Facial Movement. Ann. Otol. Rhinol. Laryngol. 109, 478–483 (2000).

20. Bègue, I. et al. Confidence of emotion expression recognition recruits brain regions outside the face perception network. Soc. Cogn. Affect. Neurosci. 14, 81–95 (2019).

21. Chen, B., Mundy, M. & Tsuchiya, N. Metacognitive Accuracy Improves With the Perceptual Learning of a Low- but Not High-Level Face Property. Front. Psychol. 10, 1712 (2019).

22. Lapate, R. C., Samaha, J., Rokers, B., Postle, B. R. & Davidson, R. J. Perceptual metacognition of human faces is causally supported by function of the lateral prefrontal cortex. *Commun*. Biol. 3, 1–10 (2020).

23. Shea, N. et al. Supra-personal cognitive control and metacognition. Trends Cogn. Sci. 18, 186–193 (2014).

24. Fuentes, C. T., Runa, C., Blanco, X. A., Orvalho, V. & Haggard, P. Does My Face FIT?: A Face Image Task Reveals Structure and Distortions of Facial Feature Representation. PLoS ONE 8, e76805 (2013).

25. Fuentes, C. T., Longo, M. R. & Haggard, P. Body image distortions in healthy adults. Acta Psychol. (Amst.) 144, 344–351 (2013).

26. Longo, M. R. & Haggard, P. An implicit body representation underlying human position sense. Proc. Natl. Acad. Sci. 107, 11727–11732 (2010).

27. Maister, L., De Beukelaer, S., Longo, M. & Tsakiris, M. The Self in the Mind’s Eye: Reverse-correlating one’s self reveals how psychological beliefs and attitudes shape our body-image. https://osf.io/f2b36 (2020) doi:10.31234/osf.io/f2b36.

28. Cunningham, D. W., Kleiner, M., Wallraven, C. & Bülthoff, H. H. Manipulating Video Sequences to Determine the Components of Conversational Facial Expressions. ACM Trans Appl Percept 2, 251–269 (2005).

29. Jeffreys, H. The Theory of Probability. (OUP Oxford, 1998).

30. Maniscalco, B. & Lau, H. A signal detection theoretic approach for estimating metacognitive sensitivity from confidence ratings. Conscious. Cogn. 21, 422–430 (2012).

31. Rouault, M., McWilliams, A., Allen, M. G. & Fleming, S. M. Human metacognition across domains: insights from individual differences and neuroimaging. Personal. Neurosci. 1, (2018).

32. Rahnev, D. et al. The Confidence Database. Nat. Hum. Behav. 4, 317–325 (2020).

33. Vickers, D. & Packer, J. Effects of alternating set for speed or accuracy on response time, accuracy and confidence in a unidimensional discrimination task. Acta Psychol. (Amst.) 50, 179–197 (1982).

34. LeDoux, J. & Bemporad, J. R. The emotional brain. J. Am. Acad. Psychoanal. 25, 525–528 (1997).

35. Stål, P., Eriksson, P.-O., Eriksson, A. & Thornell, L.-E. Enzyme-histochemical differences in fibre-type between the human major and minor zygomatic and the first dorsal interosseus muscles. Arch. Oral Biol. 32, 833–841 (1987).

36. Stål, P., Eriksson, P.-O., Eriksson, A. & Thornell, L.-E. Enzyme-histochemical and morphological characteristics of muscle fibre types in the human buccinator and orbicularis oris. Arch. Oral Biol. 35, 449–458 (1990).

37. Goodmurphy, C. W. & Ovalle, W. K. Morphological study of two human facial muscles: orbicularis oculi and corrugator supercilii. Clin. Anat. N. Y. N 12, 1–11 (1999).

38. Happak, W., Burggasser, G., Liu, J., Gruber, H. & Freilinger, G. Anatomy and Histology of the Mimic Muscles and the Supplying Facial Nerve. in The Facial Nerve (eds. Stennert, E. R., Kreutzberg, G. W., Michel, O. & Jungehülsing, M.) 85–86 (Springer, 1994). doi:10.1007/978-3-642-85090-5_23.

39. Cobo, J. L., Abbate, F., de Vicente, J. C., Cobo, J. & Vega, J. A. Searching for proprioceptors in human facial muscles. Neurosci. Lett. 640, 1–5 (2017).

40. Charles, L., Chardin, C. & Haggard, P. Evidence for metacognitive bias in perception of voluntary action. Cognition 194, 104041 (2020).

41. Arbuzova, P. et al. Measuring Metacognition of Direct and Indirect Parameters of Voluntary Movement. bioRxiv 2020.05.14.092189 (2020) doi:10.1101/2020.05.14.092189.

42. Fleming, S. M. & Lau, H. C. How to measure metacognition. Front. Hum. Neurosci. 8, 443 (2014).

43. Locke, S. M., Mamassian, P. & Landy, M. S. Performance monitoring for sensorimotor confidence: A visuomotor tracking study. Cognition 104396 (2020) doi:10.1016/j.cognition.2020.104396.

44. McIntosh, R. D., Fowler, E. A., Lyu, T. & Della Sala, S. Wise up: Clarifying the role of metacognition in the Dunning-Kruger effect. J. Exp. Psychol. Gen. 148, 1882–1897 (2019).

45. Mole, C. D., Jersakova, R., Kountouriotis, G. K., Moulin, C. J. & Wilkie, R. M. Metacognitive judgements of perceptual-motor steering performance: *Q*. J. Exp. Psychol. (2018) doi:10.1177/1747021817737496.

46. Chambon, V., Filevich, E. & Haggard, P. What is the Human Sense of Agency, and is it Metacognitive? in The Cognitive Neuroscience of Metacognition (eds. Fleming, S. M. & Frith, C. D.) 321–342 (Springer Berlin Heidelberg, 2014).

47. Froemer, R., Nassar, M. R., Stuermer, B., Sommer, W. & Yeung, N. I knew that! Confidence in outcome prediction and its impact on feedback processing and learning. BioRxiv 442822 (2018).

48. Pauen, M. Die Natur des Geistes. (S. Fischer Verlag, 2016).

49. Marcel, A. J. Agency and Self-Awareness: Issues in Philosophy and Psychology. (2003).

50. Metcalfe, J. & Greene, M. J. Metacognition of agency. J. Exp. Psychol. Gen. 136, 184–199 (2007).

51. Fourneret, P. & Jeannerod, M. Limited conscious monitoring of motor performance in normal subjects. Neuropsychologia 36, 1133–1140 (1998).

52. Mazzoni, P. & Krakauer, J. W. An Implicit Plan Overrides an Explicit Strategy during Visuomotor Adaptation. J. Neurosci. 26, 3642–3645 (2006).

53. Malone, L. A. & Bastian, A. J. Thinking About Walking: Effects of Conscious Correction Versus Distraction on Locomotor Adaptation. J. Neurophysiol. 103, 1954–1962 (2010).

54. Pauen, M. The Functional Mapping Hypothesis. Topoi 36, 107–118 (2017).

55. Chiovetto, E., Curio, C., Endres, D. & Giese, M. Perceptual integration of kinematic components in the recognition of emotional facial expressions. J. Vis. 18, 13 (2018).

56. Dobs, K., Bülthoff, I. & Schultz, J. Use and Usefulness of Dynamic Face Stimuli for Face Perception Studies—a Review of Behavioral Findings and Methodology. Front. Psychol. 9, (2018).

57. Krumhuber, E. G., Skora, L., Küster, D. & Fou, L. A Review of Dynamic Datasets for Facial Expression Research: Emot. Rev. (2016) doi:10.1177/1754073916670022.

58. Brainard, D. H. The Psychophysics Toolbox. Spat. Vis. 10, 433–436 (1997).

59. Kleiner, M. et al. What’s new in Psychtoolbox-3. Perception 36, 1–1 (2007).

60. Pelli, D. G. The VideoToolbox software for visual psychophysics: transforming numbers into movies. Spat. Vis. 10, 437–442 (1997).

61. Ekman, P. Basic emotions. Handb. Cogn. Emot. 98, 16 (1999).

62. Bagby, R. M., Parker, J. D. A. & Taylor, G. J. The twenty-item Toronto Alexithymia scale—I. Item selection and cross-validation of the factor structure. J. Psychosom. Res. 38, 23–32 (1994).

63. Lange, K., Kühn, S. & Filevich, E. “Just Another Tool for Online Studies” (JATOS): An Easy Solution for Setup and Management of Web Servers Supporting Online Studies. PLoS ONE 10, e0130834 (2015).

64. Morey, R. D., Rouder, J. N. & Jamil, T. BayesFactor: Computation of Bayes Factors for common designs. R package version 0.9. 12-4.2. Comput. Softw. Retrieved HttpsCRAN R-Proj. Orgpackage BayesFactor (2018).

65. Bulat, A. & Tzimiropoulos, G. How far are we from solving the 2D & 3D Face Alignment problem? (and a dataset of 230,000 3D facial landmarks). 2017 IEEE Int. Conf. Comput. Vis. ICCV 1021–1030 (2017) doi:10.1109/ICCV.2017.116.

66. Bürkner, P.-C. Advanced Bayesian Multilevel Modeling with the R Package brms. R J. 10, 395–411 (2018).

67. Bürkner, P.-C. brms: An R Package for Bayesian Multilevel Models Using Stan. J. Stat. Softw. 80, 1–28 (2017).

68. Dienes, Z. How Do I Know What My Theory Predicts? Adv. Methods Pract. Psychol. Sci. 2, 364–377 (2019).

69. Makowski, D. et al. bayestestR: Understand and Describe Bayesian Models and Posterior Distributions. (2020).

70. Kruschke, J. K. & Liddell, T. M. The Bayesian New Statistics: Hypothesis testing, estimation, meta-analysis, and power analysis from a Bayesian perspective. Psychon. Bull. Rev. 25, 178–206 (2018).

71. Cohen, J. Statistical power analysis for the behavioral sciences. (L. Erlbaum Associates, 1988).

72. Gelman, A., Goodrich, B., Gabry, J. & Vehtari, A. R-squared for Bayesian Regression Models. Am. Stat. 73, 307–309 (2019).

73. Doorn, J. van, Ly, A., Marsman, M. & Wagenmakers, E.-J. Bayesian rank-based hypothesis testing for the rank sum test, the signed rank test, and Spearman’s ρ. J. Appl. Stat. 47, 2984–3006 (2020).

74. JASP Team. JASP (Version 0.14)[Computer software]. JASP - Free and User-Friendly Statistical Software https://jasp-stats.org/faq/how-do-i-cite-jasp/ (2020).

75. Rahnev, D. et al. The Confidence Database. https://osf.io/h8tju (2019) doi:10.31234/osf.io/h8tju.

76. Vickers, D. & Packer, J. Effects of alternating set for speed or accuracy on response time, accuracy and confidence in a unidimensional discrimination task. Acta Psychol. (Amst.) 50, 179–197 (1982).

77. Response-Related Signals Increase Confidence But Not Metacognitive Performance | eNeuro. https://www.eneuro.org/content/7/3/ENEURO.0326-19.2020.

78. Brick, T. R., Braun, J., Harrill, C. & Yu, M. Face Modeling GUI, Version 0.2β.” Software for facial expression analysis and stimulus synthesis. (2013).

79. Surowiecki, J. The wisdom of crowds. (Anchor, 2005).

80. Fleming, S. M., Weil, R. S., Nagy, Z., Dolan, R. J. & Rees, G. Relating Introspective Accuracy to Individual Differences in Brain Structure. Science 329, 1541–1543 (2010).

